# NF-κB signaling regulates the formation of proliferating Müller glia-derived progenitor cells in the avian retina

**DOI:** 10.1101/724260

**Authors:** Isabella Palazzo, Kyle Deistler, Thanh V. Hoang, Seth Blackshaw, Andy J. Fischer

## Abstract

Neuronal regeneration in the retina is a robust, effective process in some cold-blooded vertebrates, but this process is ineffective in warm-blooded vertebrates. Understanding the mechanisms and cell-signaling pathways that restrict the reprogramming of Müller glia into proliferating neurogenic progenitors is key to harnessing the regenerative potential of the retina. Inflammation and reactive microglia are known to influence the formation of Müller glia-derived progenitor cells (MGPCs), but the mechanisms underlying this response are unknown. Using the chick retina *in vivo* as a model system, we investigate the role of the Nuclear Factor kappa B (NF-κB) signaling, a critical regulator of inflammation. We find that components of the NF-κB pathway are expressed by Müller glia and are dynamically regulated after neuronal damage or treatment with growth factors. Inhibition of NF-κB enhances, whereas activation suppresses the formation of proliferating MGPCs. Additionally, activation of NF-κB promotes glial differentiation from MGPCs in damaged retinas. With microglia ablated, the effects of NF-κB-agonists/antagonists on MGPC formation are reversed, suggesting that the context and timing of signals provided by reactive microglia influence how NF-κB-signaling impacts the reprogramming of Müller glia. We propose that NF-κB-signaling is an important signaling “hub” that suppresses the reprogramming of Müller glia into proliferating MGPCs and this “hub” coordinates signals provided by reactive microglia.

## Introduction

Müller glia are the primary type of support cell of the retina and have the capacity to reprogram into proliferating neurogenic progenitor cells (Fischer and Reh, 2001). Although Müller glia have the potential to act as a source of retinal regeneration, their regenerative potential varies greatly across vertebrate species. In the retinas of teleost fish, Müller glia readily undergo a neurogenic program to produce different types of retinal neurons and restore visual function (Bernardos et al., 2007; Fausett and Goldman, 2006; Hitchcock and Raymond, 1992; Lenkowski and Raymond, 2014). By contrast, mammalian Müller glia undergo a gliotic program after damage and fail to reprogram into proliferating progenitor-like cells (Bringmann et al., 2009; Dyer and Cepko, 2000). Interestingly, avian Müller glia have a regenerative potential that lies between that of Müller glia in fish and mammals (Fischer and Reh, 2001; Gallina et al., 2014a, 201). In the chick retina, in response to damage or exogenous growth factors, Müller glia de-differentiate, acquire a progenitor phenotype, and proliferate to produce numerous progeny; of these progeny, only a fraction differentiate into neurons (Fischer and Reh, 2001; Fischer and Reh, 2002). However, the neurogenic potential of Müller glia-derived progenitors cells (MGPCs) can be enhanced by targeting cell-signaling pathways, including Notch (Ghai et al., 2010; Hayes et al., 2007), glucocorticoid (Gallina et al., 2014b), Jak/Stat (Todd et al., 2016), and retinoic acid-signaling (Todd et al., 2018). In acutely damaged mouse retina, reprogramming of Müller glia can be driven by the forced expression of the Ascl1, a pro-neural bHLH transcription factor, and inhibition of histone de-acetylases (HDACs) (Jorstad et al., 2017; Ueki et al., 2015). Identifying pathways and factors that promote retinal regeneration in the zebrafish, but suppress regeneration in the chick and mouse, is expected to guide the development of novel therapeutic strategies that could restore vision in diseased retinas.

In damaged retinas, injured neurons and activation of immune cells are believed to initiate and guide the process of reprogramming of Müller glia into MGPCs. The immune system and pro-inflammatory signals regulate neurogenesis in zebrafish brain (Kyritsis et al., 2012). Both acute damage and chronic degeneration of the retina result in inflammation, characterized and mediated by the accumulation of reactive microglia (Graeber and Streit, 1990; Karlstetter et al., 2010; Wang and Wong, 2014). In response to damage or pro-inflammatory signals, retinal microglia rapidly respond by migrating and up-regulating the expression of cytokines (Fischer et al., 2014; Todd et al., 2019; Zelinka et al., 2012). In the chick retina, the ablation of microglia prior to damage or growth factor-treatment suppresses the formation of proliferating MGPCs (Fischer et al., 2014). Similarly, the ablation of microglia in the zebrafish retina impairs neuronal regeneration (Conedera et al., 2019; White et al., 2017). In addition, pro-inflammatory signals, such as IL6 or TNFα promote the formation of proliferating, neurogenic MGPCs in the zebrafish retina (Conner et al., 2014; Zhao et al., 2014). These pro-inflammatory cytokines are known to signal through the NF-κB pathway, among many others (Osborn et al., 1989).

NF-ĸB-signaling mediates inflammation in response to injury and infection, but also regulates cell survival, apoptosis, proliferation, and differentiation in various cellular contexts (Hayden and Ghosh, 2004). Currently, the role of NF-ĸB-signaling in the formation of MGPCs in the retina is not understood. Accordingly, in this study we investigate how NF-κB-signaling influences the reprogramming of Müller glia into proliferating MGPCs in the avian retina, and whether microglia are involved in regulating NF-κB-signaling. Our findings indicate that activation of NF-κB suppress the reprogramming of Müller glia in to proliferating MGPCs. However, this effect depends on the presence of reactive microglia. Collectively, our findings suggest that NF-κB-signaling is a key pathway in regulating the reprogramming Müller glia into proliferating progenitor cells.

## Methods and Materials

### Animals

The use of animals in these experiments was in accordance with the guidelines established by the National Institutes of Health and the Ohio State University. Newly hatched wild type leghorn chickens (*Gallus gallus domesticus*) were obtained from Meyer Hatchery (Polk, Ohio). Postnatal chicks were kept on a cycle of 12 hours light, 12 hours dark (lights on at 8:00 AM). Chicks were housed in a stainless steel brooder at 25°C and received water and Purina^tm^ chick starter *ad libitum*.

### Intraocular injections

Chickens were anesthetized via inhalation of 2.5% isoflurane in oxygen and intraocular injections performed as described previously (Fischer et al., 1999). For all experiments, the vitreous chamber of right eyes of chicks were injected with the experimental compound and the contralateral left eyes were injected with a control vehicle. Compounds were injected into chick eyes in 20 μl sterile saline with up to 30% (v/v) di-methyl-sulfoxide (DMSO) and 0.05 mg/ml bovine serum albumin added as a carrier. Compounds used in these studies included N-methyl-D-aspartate (NMDA) (9.2 or 147 μg/dose; cat# M3262; Sigma-Aldrich), FGF2 (250 ng/dose; cat# 233-FB; R&D systems), sulfasalazine (5.0 μg/dose; cat# S0883: Sigma-Aldrich), 15-Deoxy-delta12,14-prostaglandin J2 (PGJ2; 5.0 ug/dose; cat# J67427; Alfa Aesar), prostratin (5.0 μg/dose; Sigma-Aldrich), TNFSF15 (250 ng/dose; cat# RP0116C; KingFisher Biotech), DAPT (*N*-[*N*-(3,5-difluorophenacetyl-L-alanyl)]-*S*-phenylglycine *t*-butyl ester) (865 ng/dose; cat# D5942; Sigma-Aldrich), and SC75741( SC757; 2.0 μg/dose; cat# s7273; Selleck Chem). 2.0 μg of EdU (5-ethynyl-2’-deoxyuridine; cat# 900584; ThermoFisher Scientific) was injected to label proliferating cells. Injection paradigms are included in each figure.

### Fixation, sectioning and immunocytochemistry

Tissues were fixed, sectioned, and immunolabeled as described previously (Fischer et al. 2008; Fischer et al. 2009b). Working dilutions and sources of antibodies used in this study are listed in Table 1.

**Table 1:** Antibodies, sources, and working dilutions.

| Antibody | Dilution | Host | Clone/Catalog number | Source |
| --- | --- | --- | --- | --- |
| Sox2 | 1:1000 | Goat | KOY0418121 | R&D |
| Sox9 | 1:2000 | Rabbit | AB5535 | Millipore |
| CD45 | 1:200 | Mouse | HIS-C7 | Prionics |
| Pax6 | 1:1000 | Rabbit | Poly19013 | BioLegend |
| Neurofilament | 1:100 | Mouse | RT97 | DSHB |
| Pospho-Histone H3 | 1:120 | Rabbit | 06-570 | Millipore |
| Brn3 | 1:50 | Mouse | MAB1585 | Chemicon |
| HuC/D | 1:150 | Mouse | A21271 | Invitrogen |
| Glutamine Synthetase | 1:2000 | Mouse | 610517 | BD Transduction Laboratories |

None of the observed labeling was due to non-specific labeling of secondary antibodies or autofluorescence because sections labeled with secondary antibodies alone were devoid of fluorescence. Secondary antibodies included donkey-anti-goat-Alexa488/568, goat-anti-rabbit-Alexa488/568/647, goat-anti-mouse-Alexa488/568/647, goat anti-rat-Alexa488 (Life Technologies) diluted to 1:1000 in PBS plus 0.2% Triton X-100.

### Labeling for EdU

Immunolabeled tissue sections were fixed in 4% formaldehyde in PBS for 5 minutes at room temperature, washed for 5 minutes with PBS, permeabilized with 0.5% Triton X-100 in PBS for 1 minute at room temperature, and washed twice for 5 minutes in PBS. Sections were incubated for 30 minutes at room temperature in 2M Tris, 50 mM CuSO_4_, Alexa Fluor 568 or 647 Azide (ThermoFisher Scientific), and 0.5M ascorbic acid in dH_2_O. Sections were washed with PBS for 5 minutes and further processed for immunofluorescence as required.

### Terminal deoxynucleotidyl transferase dUTP nick end labeling (TUNEL)

To identify dying cells that contained fragmented DNA the TUNEL method was used. We used an *In Situ* Cell Death Kit (TMR red; cat # 12156792910; Roche Applied Science), as per the manufacturer’s instructions.

### Preparation of clodronate liposomes

The preparation of clodronate liposomes was similar to previous descriptions (Van Rooijen 1989; Zelinka et al. 2012). 50 ng cholesterol and 8 mg egg lecithin (L-α-Phosphatidyl-DL-glycerol sodium salt (Sigma P8318)) were dissolved in chloroform in a round-bottom flask. The solution was evaporated under nitrogen until a white liposome residue remained. 158 mg dichloro-methylene diphosphonate (clodronate; Sigma-Aldrich) in sterile PBS was added and mixed. Clodronate encapsulation and vesicle size normalization was facilitated by sonication at 42,000Hz for 5 mins. The liposomes were centrifuged at 10,000xg for 15 min and re-suspended in 150 ml sterile PBS. We are unable to determine the exact clodronate concentration due to the stochastic nature of the clodronate combining with the liposomes. We tittered doses to levels where >99% of microglia were ablated at 2 days after treatment.

### scRNA-seq

Retinas were acutely dissociated via papain digestion and mild trituration. Dissociated cells were loaded onto the 10X Chromium Controller using Chromium Single Cell<u>3′</u> v2 reagents. Sequencing libraries were prepared following the manufacturer’s instructions (10X Genomics), with 10 cycles used for cDNA amplification and 12 cycles for library amplification. The resulting sequencing libraries were sequenced with Paired End reads, with Read 1 (26 base pairs) and Read 2 (98 base pairs), on anNextseq500 at the Genomics Resources Core Facility (High Throughput Center) at Johns Hopkins University. Raw sequence data was processed with Cell Ranger software (10X Genomics) to align sequences, de-multiplex, annotated to ENSMBL databases, count reads, assess levels of expression and construct gene-cell matrices. t-distributed stochastic neighbor embedding (tSNE) plots were generated and probed using Cell Ranger and Cell Browser software (10X Genomics). The tSNE plots were generated via aggregate cluster analysis of 9 separate cDNA libraries, including 2 replicates of control undamaged retinas, and retinas at different times after NMDA-treatment. The identity of clustered cells was established using known cell-type specific markers. Seurat was used to generate violin/scatter plots for candidate genes in identified clusters of cells (Powers and Satija 2015; Satija et al. 2015). Monocle 2.1 was used to establish pseudotime trajectories and levels of gene expression across pseudotime for cells identified as Müler glia or MGPCs (Trapnell et al. 2014). Sc-RNA seq libraries are available: https://proteinpaint.stjude.org/F/2019.retina.scRNA.html

### Photography, measurements, cell counts and statistics

Wide-field photomicroscopy was performed using a Leica DM5000B microscope equipped with epifluorescence and Leica DC500 digital camera or Zeiss AxioImager M2 equipped with epifluorescence and Zeiss AxioCam MRc. Confocal images were obtained using a Leica SP8 imaging system at the Department of Neuroscience Imaging Facility at The Ohio State University. Images were optimized for color, brightness and contrast, multiple channels overlaid, and figures constructed by using Adobe Photoshop. Cell counts were performed on representative images. To avoid the possibility of region-specific differences within the retina, cell counts were consistently made from the same region of retina for each data set.

Similar to previous reports (Fischer et al. 2009a; Fischer et al. 2009b; Fischer et al. 2010; Ghai et al. 2009), immunofluorescence was quantified by using ImagePro6.2 (Media Cybernetics, Bethesda, MD, USA). Identical illumination, microscope, and camera settings were used to obtain images for quantification. Retinal areas were sampled from 5.4 MP digital images. These areas were randomly sampled over the inner nuclear layer (INL) where the nuclei of the bipolar and amacrine neurons were observed. Measurements of immunofluorescence were performed using ImagePro 6.2 as described previously (Ghai et al., 2009; Stanke et al 2010; Todd and Fischer 2015). The density sum was calculated as the total of pixel values for all pixels within thresholded regions. The mean density sum was calculated for the pixels within threshold regions from ≥5 retinas for each experimental condition. GraphPad Prism 6 was used for statistical analyses.

Central retina was defined as the region within a 3mm radius of the posterior pole of the eye, and peripheral retina was defined as an annular region between 3mm and 0.5mm from the CMZ. The identity of EdU-labeled cells was determined based on previous findings that 100% of the proliferating cells in the chick retina are comprised of Sox2/9^+^ Müller glia in the INL/ONL, Sox2/9/Nkx2.2^+^ Non-astrocytic Inner Retinal Glial (NIRG) cells in the IPL, GCL, and NFL (the NIRG cells do not migrate distally into the retina), and CD45^+^ microglia (Fischer et al. 2010; Zelinka et al. 2012). Sox2^+^ nuclei in the INL were identified as Müller glia based on their large size and fusiform shape which was distinctly different from the Sox2^+^ nuclei of cholinergic amacrine cells which are small and round (Fischer et al. 2010).

GraphPad Prism 6 was used for statistical analyses and generation of histograms and bar graphs. Where statistical significance of difference was determined between treatment groups accounting for intra-individual variability within a biological sample, we performed a two-tailed, paired t-test. Where significance of difference was determined between two treatment groups comparing inter-individual variability we performed two-tailed, unpaired t-tests. When evaluating significance in difference between multiple groups we performed ANOVA followed by Tukey’s Test.

## Results

### Expression patterns of NF-κB signaling components in damaged retinas

Previous studies have indicated that NMDA-induced excitotoxic damage in the mouse retina results in the activation of NF-ĸB in Müller glia (Lebrun-Julien et al., 2009). We failed to identify antibodies to components of the NF-κB pathway, including RelA/p65, phospho-p65, phospho-IKBα/β, or RelB, that produced robust and reproducible patterns of labeling. Accordingly, we queried single cell RNA-sequencing (scRNA-seq) databases generated from chick retinas at different times after NMDA-treatment. Unbiased tSNE plots of aggregate scRNA-seq databases revealed discrete clustering of Müller glia from control retinas and at 24 hrs after NMDA-treatment, whereas Müller glia from 48 and 72 hrs after treatment clustered together (Fig. 1a). Müller glia were identified based on combined expression of *Vim, Slc3a1, Glul* and *Rlbp1*, and MGPCs were identified based on down-regulation of *Glul* and *Rlbp1* and up-regulation of *Pcna, Cdk1* and *Top2a* (Fig. 1b). Microglia were identified based on the distinct expression of *C1qa, C1qb, C1qc, Csf1r, C3ar1, Ccl4* and *Hhex* (Fig. 1c).

**Figure 1:**
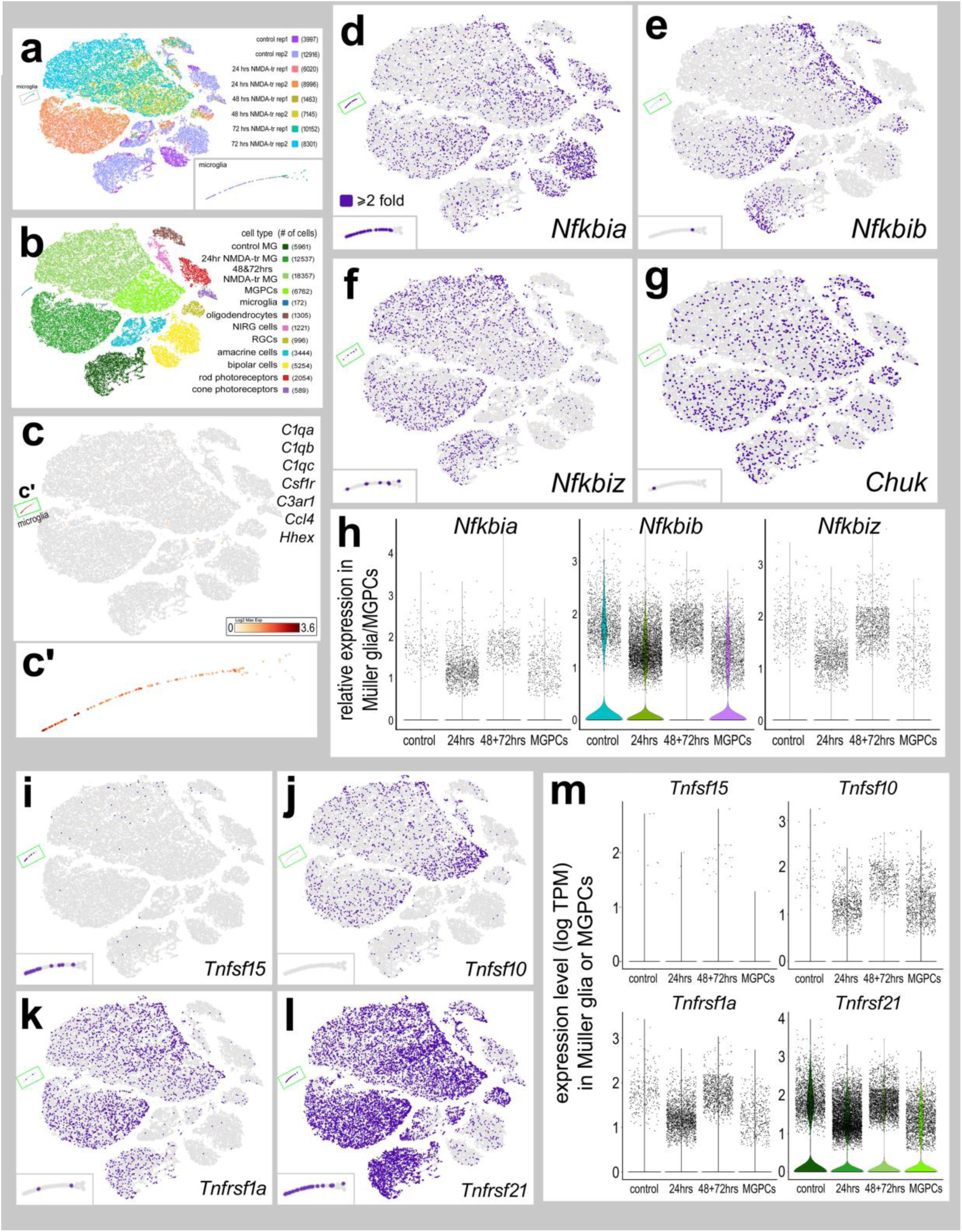
Expression patterns of NF-κB signaling components in damaged retinas: Libraries for scRNA-seq were established for cells from control retinas and from retinas at different times after NMDA-treatment. Each dot represents one cell. Cells were sampled from control retinas (rep1 3997 cells, rep2 12916 cells), and from retinas at 24hrs (rep1 6020 cells, rep2 8996 cells), 48hrs (rep1 1463 cells, rep2 7145 cells), and 72hrs (rep1 10152 cells, rep2 8301 cells) after NMDA-treatment (**a**). tSNE plots show distinct clustering of different retinal cell populations and numbers of cells surveyed within each cluster (in parentheses) (**b**). Microglia were identified based on expression *of C1qa, C1qb, C1qc, Csf1r, C3ar1, Ccl4, and Hhex* **(c,c’)**. Müller glia were identified based on significant (≥8-fold; purple dots) collective expression of *Vim*, *Glul*, *Rlbp1,* and *Slc1a3*. t-SNE plots for expression (≥2-fold; purple dots) of *Nfkbia, Nfkbib, Nfkbiz, Chuk, Tnfsf15, Tnfsf10, Tnfrsf1a* and *Tnfrsf21* (**d**-**l**). Violin/scatter plots illustrate expression levels of *Tnfsf15, Tnfsf10, Tnfrsf1a* and *Tnfrsf21* in Müller glia and MGPCs in controls and at different times after NMDA-treatment (**m**).

We probed for different components of the NF-κB pathway, including *Nfkbia, Nfkbib, Nfkbiz* and *Chuk*. NF-κB transcription factors include P65 (RelA), RelB, c-Rel, P50 (*Nfkb1*), and P52 (*Nfkb2*) (Zhang et al., 2017). NF-κB signaling activity is regulated by cytoplasmic Inhibitor of kappa B (IκB), which is comprised of IkBα (*Nfkbia*), IκBβ (*Nfkbib*), IκBε (*Nfkbie*), IκBγ (*Ikbkg)*, IkBζ (*Nfkbiz*), which mask the nuclear localization sequences of NF-κB transcription factors (Ghosh et al., 1998). Inhibitor of kappa B Kinases (IKKs; *Chuk* and *Ikbkb*) phosphorylate IkB, targeting it for ubiquitination and degradation, thereby resulting in liberation of NF-κB transcription factors (Karin and Ben-Neriah, 2000; Zhang et al., 2017). *Nfkbia* was prominently expressed in microglia, bipolar cells and NIRG cells, and had scattered expression, at relatively high levels, in Müller glia, MGPCs, amacrine cells and photoreceptors (Fig. 1d,h). Among Müller glia, number of Nfkbia-expressing cells were most abundant at 24 hrs after NMDA-treatment (Fig. 1h). By comparison, *Nfkbib* was predominantly expressed by Müller glia in normal and damaged retinas, and by MGPCs, but was less abundant in Müller glia at 48 and 72 hrs after NMDA-treatment (Fig. 1e,h). Similarly, *Nfkbiz* was predominantly expressed by Müller glia in control and damaged retinas, and was less abundant in MGPCs (Fig. 1f,h). *Chuk* was widely expressed by Müller glia, MGPCs, NIRG cells, oligodendrocytes, amacrine cells, bipolar cells and ganglion cells (Fig. 1g).

TNF-related ligands are known to activate NF-κB-signaling in different cellular contexts (Hayden and Ghosh, 2014; Osborn et al., 1989; Schütze et al., 1992; Schütze et al., 1995). Accordingly, we probed for TNF-related ligands and receptors in our scRNA-seq libraries. TNFα has not been identified in the chick genome, but it has been suggested that chicken Tumor Necrosis Factor Super Family 15 (TNFSF15)/TL1A may function in its place (Migone et al., 2002; Takimoto et al., 2005). We found that *Tnfsf15* was only detected in microglia (Fig. 1i), consistent with scRNA-seq data from the mouse retina where microglia are the only source of *Il1a, Il1b* and *Tnf* (Todd et al., 2019). By comparison, *Tnfsf10* was detected in relatively few Müller glia in control retinas, but was expressed in many Müller glia in damaged retinas and in MGPCs (Fig. 1j,m). Other isoforms of TNF-related ligands, including *Tnfsf6, Tnfsf8, and Tnfsf11* were not expressed at significant levels (not shown). We detected expression of TNFSF receptors predominantly in Müller glia in control and damaged retinas. In control retinas, *Tnfrsf1a* was detected at relatively high levels in scattered Müller glia (Fig. 1k,m). In damaged retinas, *Tnfrsf1a* was detected in many Müller glia at 24hrs, in decreased numbers at 48 and 72hrs after NMDA-treatment, and relatively few MGPCs (Fig. 1k,m). By comparison, the expression of *Tnfrsf21* was widespread in control Müller glia, in Müller glia from damaged retinas, and in MGPCs (Fig. 1l,m). TNFSF15 is an orthologue to mammalian TNFα and is similar in sequence to human TL1a (Takimoto et al., 2005).

To better understand the expression patterns of NF-κB-related genes in Müller glia and MGPCs, we assessed relative levels of expression across pseudotime. Using Monocle (Trapnell et al., 2014), pseudotime states and trajectories were established by comparing the highly variable genes among Müller glia and MGPCs. This analysis revealed 5 distinct pseudotime states and the trajectories include distinct branches for resting Müller glia, activated Müller glia, transitional Müller glia, and MGPCs (Fig. 2a). Resting Müller glia from control retinas were located to the far left of the pseudotime axis, and expressed high levels of mature glial markers, such as *Glul* (Fig. 2b-c). MGPCs from 48 and 72 hrs after NMDA-treatment were located to the far right of the pseudotime axis and expressed high levels of markers for proliferation and progenitor cells, such as *Cdk1* (Fig. 2b-c). Expression across pseudotime revealed small decreases in relative levels *Nfkbiz* (Fig. 2d), whereas levels of *Nfkbia* were unchanged and high-expressing Muller glia were scattered evenly across pseudotime (Fig. 2d). By comparison, levels of *Nfkbib* decreased in transitional glia and then increased toward MGPCs (Fig. 2d). Expression of *Tnfrsf10* increased over pseudotime toward MGPCs, whereas *Tnfrsf1a* was slightly increased in transitional glia and decreased in MGPCs (Fig. 2d). By comparison, expression of *Tnfrsf21* was relatively high in resting MG, low in transitional MG, elevated in activated MG, and decreased in MGPCs (Fig. 2d). Taken together, these findings indicate that essential components of NF-κB signaling are dynamically expressed in Müller glia after damage and during the process of reprogramming into MGPCs.

**Figure 2:**
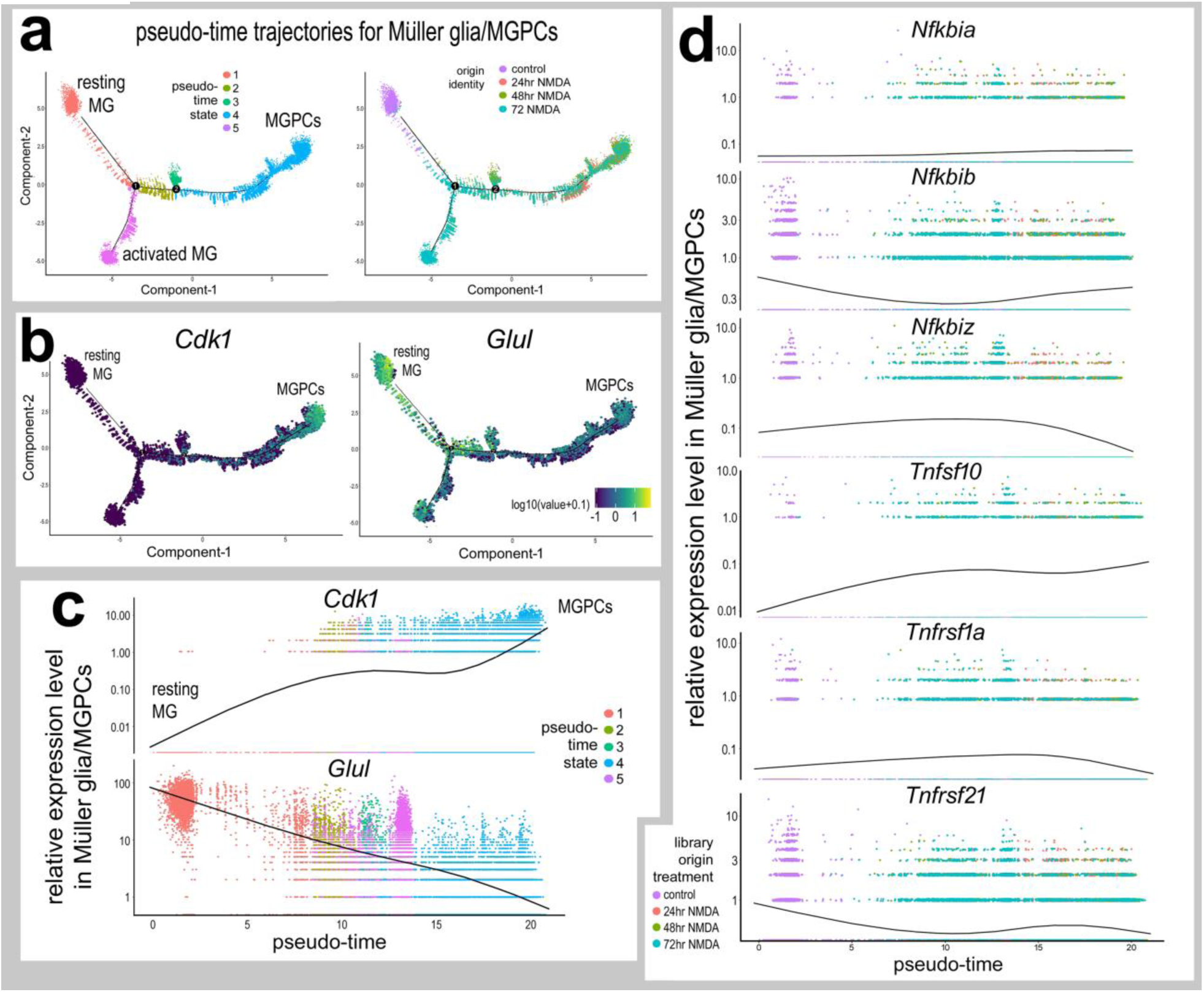
Pseudotime analysis of NF-κB-related genes in Müller glia and MGPCs. Seurat objects identified as Müller glia and MGPCs were re-embedded for pseudotime analysis. Monocle was used to generate pseudotime trajectories in an unbiased manner guided by the differentially expressed genes. Pseudotime states included: (1; peach) resting Müller glia, (2; olive) transitional MG, (3; green) transitional MG, (4; blue) MGPCs, and (5; magenta) activated Müller glia (**a**). Expression of *Glul* and *Cdk1* are shown along pseudotime (**b**). Reduction of pseudotime to the X-axis placed the resting Müller glia to the far left and the MGPCs to the far right (**c**). Relative levels of expression among Müller glia and MGPCs across pseudotime was assessed for components of the NF-κB-pathway*, Nfkbia, Nfkbib*, *Nfkbiz,* and TNF-related factors and receptors *Tnfsf10, Tnfrsf1a, Tnfrsf21* (**d**).

### NF-κB-signaling regulates the formation of MGPCs in damaged retinas

To determine whether NF-ĸB-signaling influences the formation of MGPCs in the chick retina, we applied small molecule activators/inhibitors of NF-ĸB-signaling to NMDA-damaged retinas. We tested whether small molecule antagonists, sulfasalazine, 15-deoxy-delta-12,14-prostaglandin J2 (PGJ2), or SC75741(SC757), that act at different levels of the NF-ĸB pathway, influence the formation of proliferating MGPCs Sulfasalazine is an anti-inflammatory agent that potently suppresses NF-κB activity by preventing IκBα phosphorylation and degradation, resulting in persistent sequestration of NF-κB transcription factors in the cytoplasm (Wahl et al., 1998). Sulfasalazine has also been shown to act at the cysteine/glutamate-antiporter (*Slc7a11*) (Gout et al., 2001), but this transporter is not expressed at detectable levels in scRNA-seq libraries (not shown). PGJ2 is a derivative of prostaglandin D2, and acts as a ligand for peroxisome proliferator-activated receptor γ (PPARγ) to regulate inflammation (Ricote et al., 1998; Straus et al., 2000), but also acts to inhibit NF-κB signaling in a PPARγ-independent manner (Lindström and Bennett, 2005). PGJ2 represses IKK activity, thus reducing IκBα phosphorylation degradation (Lindström and Bennett, 2005; Straus et al., 2000). Additionally, PGJ2 covalently interacts with P50, an NF-κB transcription factor, to directly block its DNA-binding ability (Cernuda-Morollón et al., 2001; Straus et al., 2000). SC757 belongs to a novel class of NF-κB inhibitors (Leban et al., 2007) that acts by blocking DNA-binding of P65, without affecting nuclear translocation or IκBα (Ehrhardt et al., 2013).

Application of NF-κB inhibitors after a low dose of NMDA (63 nmol) significantly increased numbers of proliferating MGPCs. Treatment with sulfasalazine, PGJ2, or SC757 following NMDA significantly increased numbers of Sox9^+^/EdU^+^ cells (Figs. 3a-d) and numbers of pHisH3^+^/neurofilament^+^ mitotic cells in the INL and ONL (Figs. 3e-g). In addition, application of sulfasalazine following NMDA resulted in increased expression of stem cell associated transcription factor Pax6 in Sox2-positive cells (Fig. 3h-j), suggesting that inhibition of NF-κB promotes the reprogramming of Müller glia into progenitor-like cells. Treatment of NMDA-damaged retinas with PGJ2 or SC757, but not sulfasalazine, resulted in decreased proliferation of microglia (Supplemental Fig. 1). Collectively, these findings indicate that inhibition of NF-κB at different levels of the pathway promotes the formation of proliferating MGPCs in damaged retinas.

**Figure 3:**
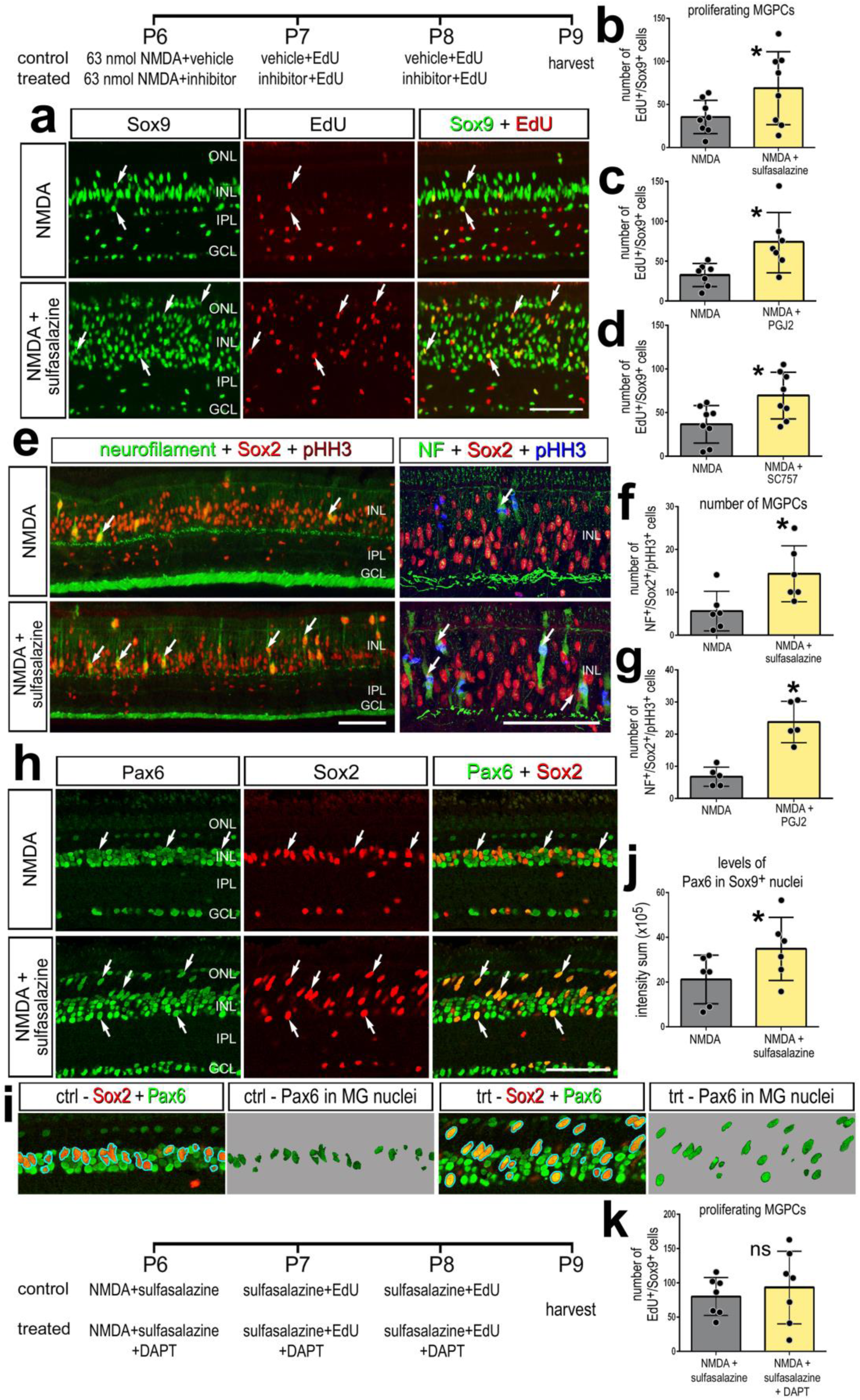
Inhibition of NF-κB-signaling promotes MGPC proliferation after NMDA damage. NMDA-damaged retinas were treated with 3 consecutive daily doses of sulfasalazine, PGJ2, or SC757, or vehicle controls. Retinas were harvested 72 hrs after damage (**a-j**). Retinal sections were labeled with Sox9 (green; **a**), EdU (red; **a**), neurofilament (green; **e**), Sox2 (red; **e,h-i**), pHistone H3 (blue; **e**), or Pax6 (green; **h-i**). NMDA-damaged retinas were treated with 3 consecutive daily doses of sulfasalazine with or without the Notch inhibitor DAPT, and retinas were harvested 72 hrs after damage. The histograms in **b**, **c**, **d, f, g,** and **k** illustrate the mean (± SD and individual data points) of proliferating cells (n≥6) and histogram in **j** represents mean (± SD and individual data points) pixel intensity above threshold for Pax6 immunofluorescence. Significance of difference (*p<0.05) was determined by using a paired *t*-test. Arrows indicate proliferating MGPCs in **a**. Arrows in **d** represent mitotic figures. Arrows in **h** represent MGPCs expressing Pax6. The calibration bars in panels **a, e,** and **h** represents 50 µm. Abbreviations: ONL – outer nuclear layer, INL – inner nuclear layer, IPL – inner plexiform layer, GCL – ganglion cell layer.

To understand how NF-κB signaling fits into the network of cell-signaling pathways that regulate the formation of MGPCs we investigated interaction with Notch-signaling. In the chick retina, Notch-signaling is required for the proliferation of MGPCs; gamma-secretase inhibitor, DAPT, suppresses MGPC-proliferation, but enhances neuronal differentiation (Ghai et al., 2010; Hayes et al., 2007). To investigate whether NF-κB coordinates with Notch-signaling we treated damaged retinas with sulfazalasine and DAPT. There was no significant difference in the number of Sox2^+^/EdU^+^ cells in damaged retinas treated with sulfasalazine and DAPT compared to damaged retinas treated with sulfasalazine alone (Fig. 3k). This finding suggests that NF-κB takes presidence or acts downstream of Notch to regulate the proliferation of MGPCs.

We next investigated whether activation of NF-κB following damage influenced the formation of MGPCs. Prostratin activates NF-κB signaling by stimulating phosphorylation and degradation of IκBα in an IKK-dependent manner (Williams et al., 2004). Prostratin-induced NF-κB activation is likely mediated by PKC (Lin et al., 2000; Williams et al., 2004). Additionally, we applied TNFSF15 to damaged retinas. TNF-ligands are known to activate NF-κB-signaling (Hayden and Ghosh, 2014; Schütze et al., 1995). TNFSF15/TL1a activates NF-κB through TNFRSF25/DR3 and also binds TNFRSF21/DR6 (Migone et al., 2002). Orthologues for mammalian TNFα and TNFRSF25 have not been identified in the chick genome. Chicken TNFSF15 is homologous to mammalian TL1a, and it is likely that TL1a/TNFSF15 acts as TNFα in the chick (Takimoto et al., 2005).

We found that treatment with prostratin after a high dose of NMDA (1 µmol) resulted in a significant decrease in numbers of Sox2^+^/EdU^+^ cells in the INL (Fig. 4a-b), while there was no significant change in the proliferation of microglia (Fig. 4d). Additionally, there was a significant decrease in the number of Sox2^+^/NF^+^/pHisH3^+^ mitotic cells in the INL/ONL of NMDA damaged retinas treated with prostratin relative to NMDA controls (Fig. 4 e-f). Treatment with TNFSF15 following NMDA did not influence the proliferation of MGPCs; there was no significant difference in numbers of Sox9^+^/EdU^+^ cells between control and treated retinas (Fig. 4c). It is possible that levels of TNF-ligands produced by microglia were saturated in damaged retinas and, thus, addition of exogenous TNFSF15 had no significant effect. However, TNFSF15-treatment recruited microglia to the vitread surface of the retina in control and NMDA-damaged retinas (Supplemental Figure 2), suggesting a chemotactic influence upon the immune cells.

**Figure 4:**
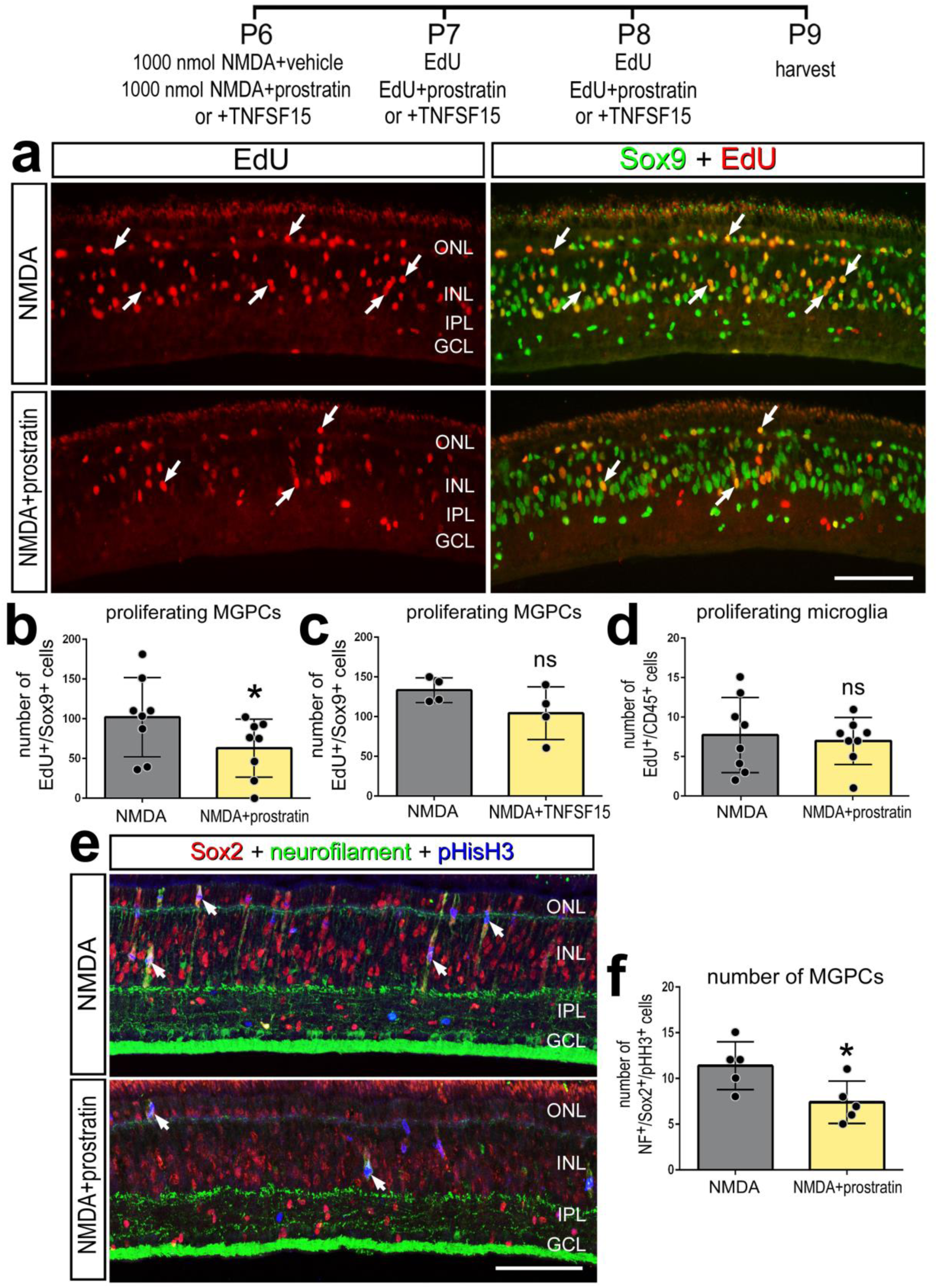
Stimulation of NF-κB signaling suppresses MGPC proliferation in damaged retinas. NMDA-damaged retinas were treated with 3 consecutive daily doses of prostratin or TNFSF15 or vehicle controls, and retinas were harvested 72 hrs after damage. Retinal sections were labeled with Sox9 (green; **a**), EdU (red; **a**), neurofilament (green; **e**), Sox2 (red; **e**), or pHistone H3 (blue; **e**). The histograms in **b**, **c**, **d** and **f** illustrate the mean (± SD and individual data points) of proliferating cells. Significance of difference (*p<0.05) was determined by using a paired *t*-test. Arrows indicate proliferating MGPCs in **a**. The calibration bars in panels **a** and **e** represent 50 µm. Abbreviations: ONL – outer nuclear layer, INL – inner nuclear layer, IPL – inner plexiform layer, GCL – ganglion cell layer.

### Inhibition of NF-κB is neuroprotective against NMDA-induced retinal damage

Levels of retinal damage and cell death are known to positively correlate with the formation of MGPCs (Fischer and Reh, 2001; Fischer and Reh, 2003), NF-κB is known to influence inflammation and neuronal survival (Hayden and Ghosh, 2008; Lanzillotta et al., 2015; Lebrun-Julien et al., 2009; Schneider et al., 1999). Thus, we investigated whether inhibition of NF-κB influenced cell death and neuronal survival, which may secondarily impact the formation of MGPCs. In the chick retina, NMDA induces cell death within 4 hrs, with numbers of dying cells peaking around 24 hrs, and continuing through to 72 hrs after treatment (Fischer et al., 1998; Fischer et al., 2015). In retinas treated with sulfasalazine following NMDA (1 µmol or 63 nmol), we found significantly fewer TUNEL-positive cells at 4 hrs, 24hrs, and 72hrs after damage relative to NMDA alone (Fig. 5a-d). Consistent with these findings, numbers of TUNEL-positive cells were significantly reduced in NMDA-damaged retinas treated with PGJ2 compared to controls (Fig. 5e). By comparison, application of prostratin following a NMDA (1 µmol) had no effect upon numbers of dying cells (Fig. 4f), whereas application of TNFSF15 following NMDA resulted in a modest, but significant, increase in numbers of dying cells (Fig. 4g).

**Figure 5:**
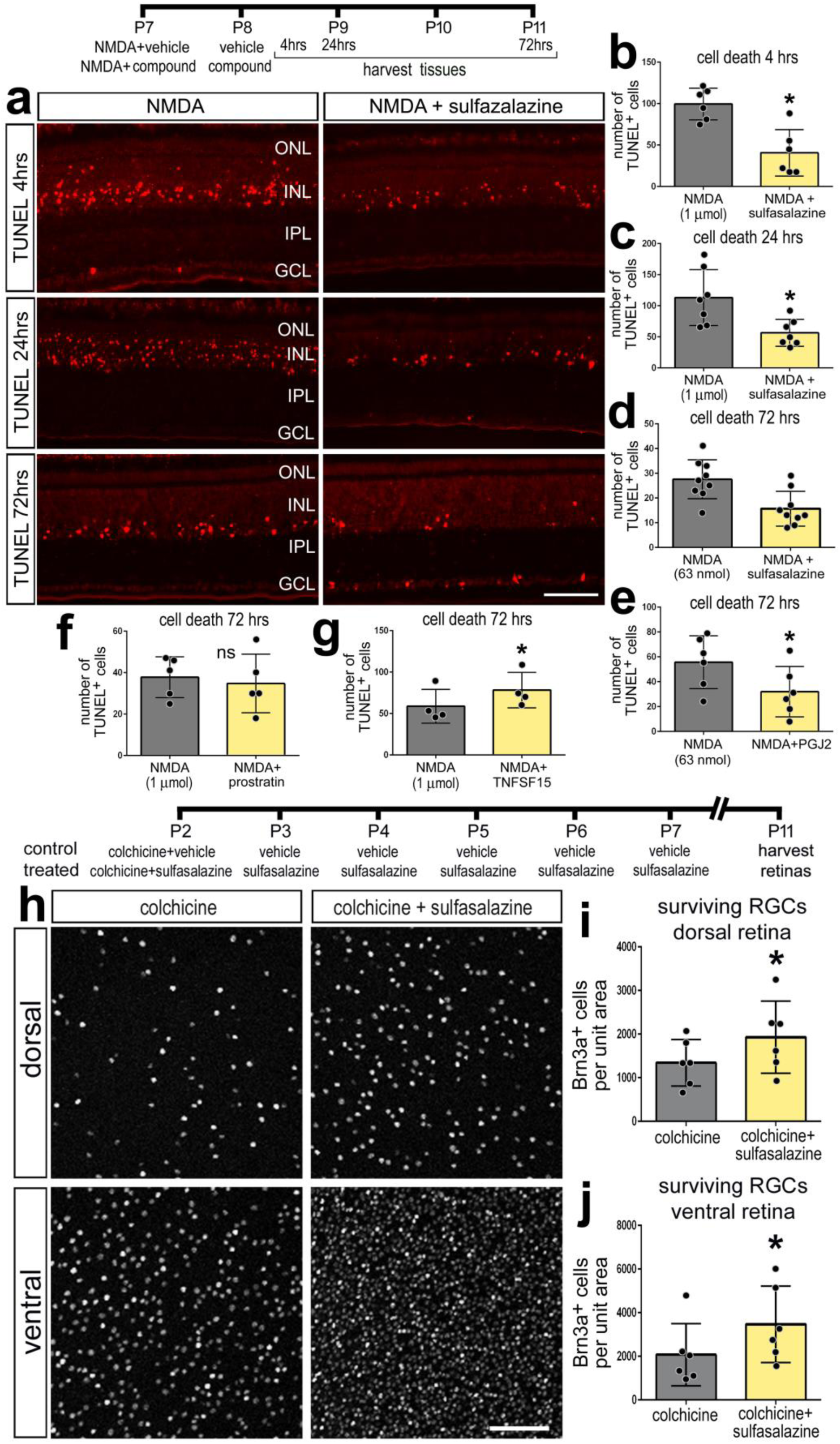
Inhibition of NF-κB is neuroprotective to different types of retinal neurons. NF-κB inhibitors (sulfasalazine or PGJ12) or activators (prostratin or TNFSF15) were applied after NMDA- or colchicine-induced damage. Retinas harvested at different times after damage. Retinas were harvested at 4 hrs (**a,b**), 24 hrs (**a**,**c**), or 72 hrs (**a**,**d-g**). In retinal sections, dying cells containing fragmented DNA were labeled with TUNEL (**a**). Histograms in **b-g** represent the mean number (± SD and individual data points) of TUNEL-positive cells. Colchicine-damaged retinas were treated with sulfasalazine or vehicle control, followed 6 consecutive daily treatments of sulfasalazine or vehicle, and retinas were harvested 9 days after colchicine treatment (**h-j**). Retinal whole-mounts were labeled with antibodies to Brn3 (**h**). Histograms represent the mean number (± SD and individual data points) of ganglion cells in dorsal (**i**) and ventral (**j**) regions of the retina (n≥6 animals). Significance of difference (*p<0.05) was determined by using a paired *t*-test. The calibration bars in panels **a** and **h** represent 50 µm. Abbreviations: ONL – outer nuclear layer, INL – inner nuclear layer, IPL – inner plexiform layer, GCL – ganglion cell layer.

We next examined whether NF-κB-signaling influences the survival of retinal ganglion cells. NMDA damage primarily destroys amacrine cells, bipolar cells, and horizontal cells in the chick retina, whereas colchicine treatment causes ganglion cell death within 3 days of treatment in newly hatched chicks (Fischer et al., 1998; Stanke and Fischer, 2010). Application of sulfasalazine following colchicine treatment resulted in significantly greater numbers of surviving Brn3^+^ ganglion cells in both dorsal and ventral regions relative to controls (Fig. 5h-j). Taken together, these data indicate that TNFSF15 and NF-κB signaling promotes the death of retinal neurons, while inhibition of NF-κB promotes neuronal survival.

### NF-κB-signaling promotes glial cell-fate after damage

During neural development NF-κB-signaling acts as a pro-gliogenic pathway (Fujita et al., 2011; Keohane et al., 2010; Mondal et al., 2004). Thus, we tested whether inhibition of NF-κB with sulfasalazine following NMDA-damage influenced the differentiation of newly generated cells. We found a significant increase in the number of newly generated cells, but failed to find a significant difference in the percentage of progeny that differentiated into neurons (EdU^+^/HuD^+^; Fig 6c or EdU^+^/Otx2^+^; not shown), or the percentage of progeny that differentiated as glia (EdU^+^/GS^+^) (Fig. 6a-c). We next tested whether activation of NF-κB influenced cellular differentiation. NF-κB inhibitor (sulfasalazine) was applied with NMDA at P6, and at P7 and P8, to increase numbers of proliferating MGPCs. Then, NF-κB activator (prostratin) was applied at P11 and P112, when MGPCs have undergone proliferation, and retinas were harvested at P15. We found no significant difference in the number of newly generated EdU^+^ cells between treated and control conditions (Fig. 6e). However, we found a significant 2-fold increase in the percentage of MGPC-progeny that differentiated into glia (EdU^+^/GS^+^) (Fig. 6d-f). There was no significant difference in the differentiation of newly generated neurons (EdU^+^/HuD^+^; Fig 6f, or EdU^+^/Otx2^+^; not shown). These data indicate that activation of NF-κB promotes glial cell fate from the progeny of MGPCs.

**Figure 6:**
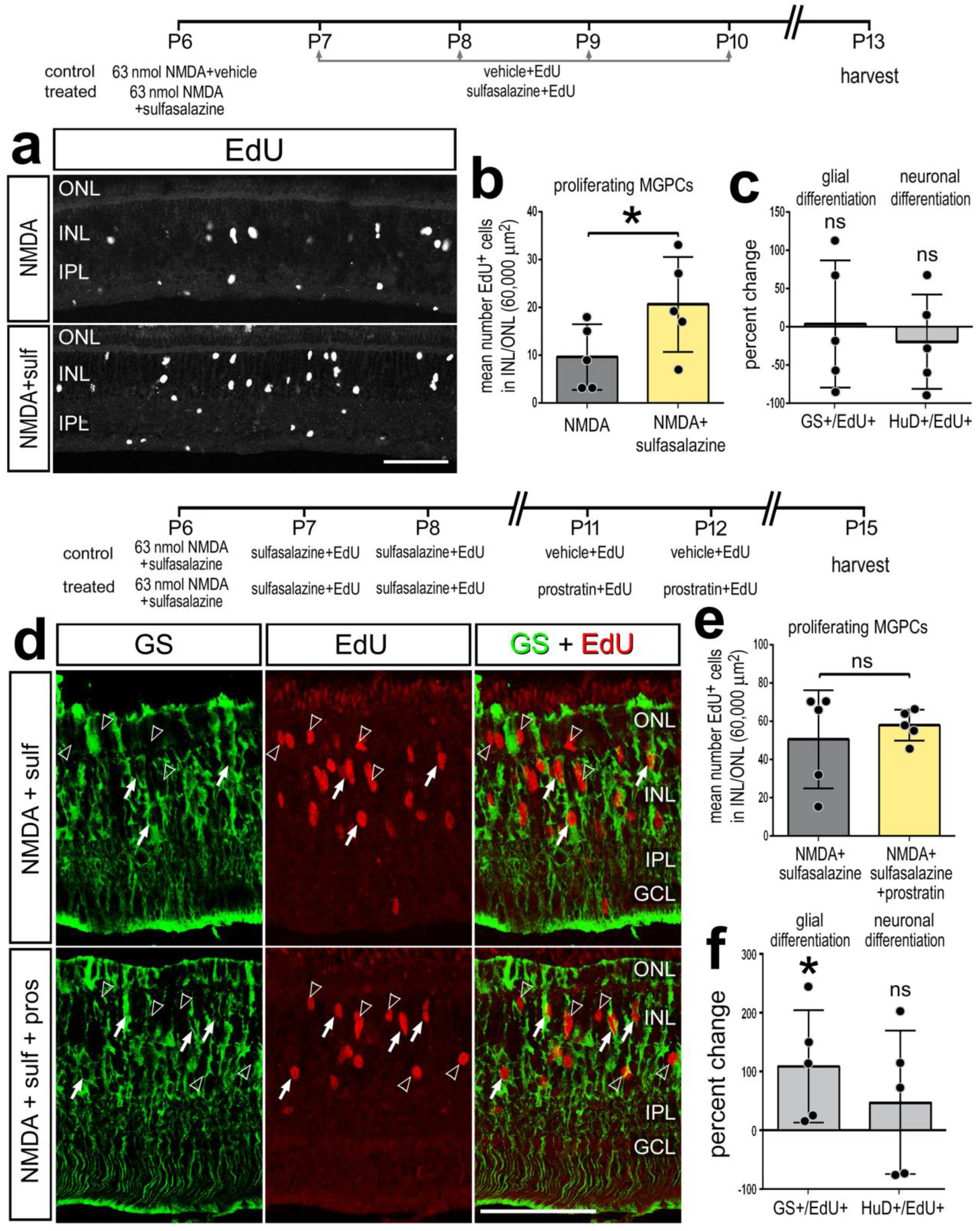
NF-κB signaling promotes glial cell-fate after damage. Retinas were treated with NMDA and vehicle (control) or NMDA and sulfasalazine (treated) at P6, followed by vehicle or sulfasalazine at P7, P8, P9, P10, and retinas harvested at P13 (**a-c**). Alternatively, retinas were treated with NMDA and vehicle (control) or NMDA and sulfasalazine (treated) at P6, followed by vehicle or sulfasalazine at P7 and P8, followed by vehicle or prostratin at P11 and P12, and retinas harvested at P15 (**d-f**). Retinal sections were labeled for EdU (**a**; grayscale and **d**; red) and GS (**d**; green). Histograms in **b** and **e** represent the mean number (± SD and individual data points) of proliferating MGPCs. Histograms in **c** and **f** represent the mean percentage change (± SD and individual data points) of EdU-positive cells that are co-labeled for GS (Müller glia) or HuD (neurons). Significance of difference (*p<0.05) was determined by using a paired *t*-test. Calibration bars in **a** and **d** represent 50 µm. Open arrows indicate proliferating MGPCs, closed arrows indicate newly generated GS-positive cells (**d**). Abbreviations: ONL – outer nuclear layer, INL – inner nuclear layer, IPL – inner plexiform layer, GCL – ganglion cell layer.

### The impact of NF-κB-signaling on MGPC proliferation after damage is dependent on the presence of reactive microglia

Signals from reactive microglia are known to impact the reprogramming of Müller glia into neurogenic MGPCs (Conner et al., 2014; Fischer et al., 2014; Nelson et al., 2013; White et al., 2017; Zhao et al., 2014). We performed scRNA-seq on normal and damaged retinas with and without microglia. We ablated retinal microglia via an intraocular injection of clodronate liposomes, which ablates >99% of microglia within 2 days of treatment (Zelinka et al., 2012). Retinas (± microglia) were harvested for scRNA-seq at 24 hrs after treatment with saline or NMDA, objects identified as Müller glia were isolated, and analyses were performed. Regardless of the presence of microglia, unbiased tSNE plots revealed distinct clustering of Müller glia from saline-treated retinas and clustering of Müller glia from damaged retinas (Fig. 7a). Relative expression levels of *Vim* were not significantly different in the absence of microglia (Figs. 7b,c), whereas damage-induced down-regulation of *Rlbp1* was not as pronounced in the absence of microglia (Figs. 7b,c). Although the absence of microglia had no effects upon the relative numbers of Müller glia expressing NF-κB components in saline-treated retinas, numbers of Müller glia expressing *Nfkbia*, *Nfkbib* and *Tnfrsf21* were reduced in NMDA-damaged retinas missing reactive (Fig. 7c). These findings suggest that the NF-κB pathway may be diminished in Müller glia from damaged retinas devoid of reactive microglia.

**Figure 7:**
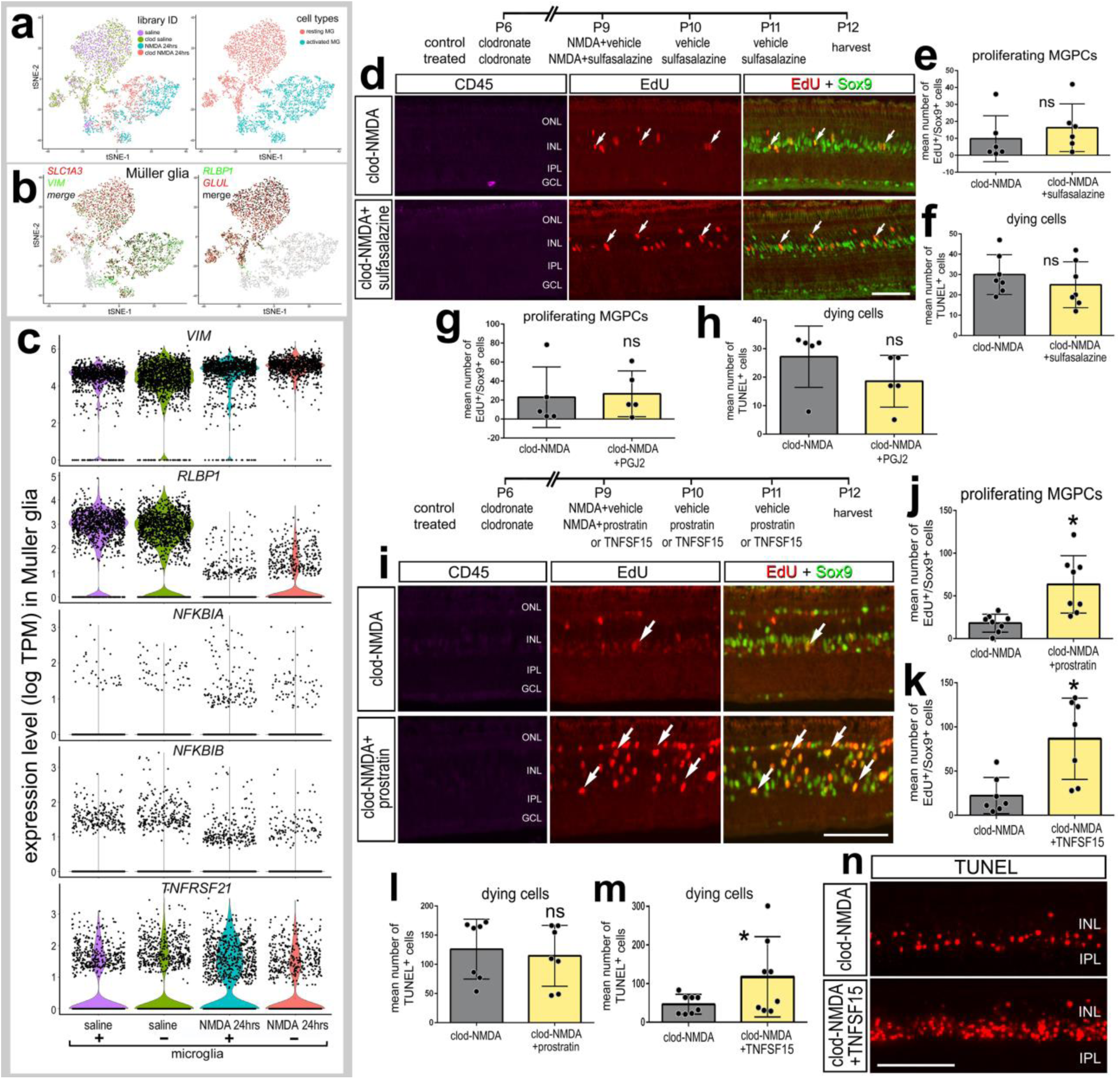
The impact of NF-κB-signaling on MGPC proliferation depends on the presence of reactive microglia. Libraries for scRNA-seq were established for cells from retinas 24 hrs after saline or NMDA-treatment with (control) and without microglia (clodronate liposomes). Müller glia were bioinformatically isolated from saline-treated retinas with microglia (1107 cells) and without microglia (1462 cells), and from NMDA-damaged retinas with microglia (1107 cells) and without microglia (1009 cells) (**a**). tSNE plots show distinct clustering of Muller glia from saline and NMDA-damaged retinas (**a**). Müller glia were identified based on collective expression of *Vim*, *Glul*, *Rlbp1,* and *Slc1a3* (**b**). Violin/scatter plots illustrate expression levels of *Vim, Rlbp1, Nfkbia, Nfkbib* and *Tnfrsf21* in Müller glia controls (± microglia) and at 24 hrs after NMDA-treatment (± microglia) (**c**). Retinas were treated with clodronate-liposomes at P6, NMDA plus vehicle or sulfasalazine at P9, vehicle or sulfasalazine plus EdU at P10 and P11, and retinas harvested at P12 (**d**-**h**). Alternatively, retinas were treated with clodronate-liposomes at P6, NMDA plus vehicle or prostratin or TNFSF15 at P9, EdU plus vehicle or prostratin or TNFSF15 at P10 and P11, and retinas harvested at P12 (**i**-**n**). Retinal sections were labeled for EdU (red; **d,i**), Sox9 (green; **d,i**), CD45 (magenta; **d**,**i**), or TUNEL (**n**). Histograms in **e, g, j** and **k** represent the mean number (± SD and individual data points) of proliferating MGPCs. Histograms in **f, h, l** and **m** represent the mean number (± SD and individual data points) of TUNEL-positive cells. Arrows in **d** and **i** represent proliferating MGPCs. Significance of difference (*p<0.05) was determined by using a paired *t*-test. Calibration bars in **d, i,** and **n** represent 50 µm. Abbreviations: ONL – outer nuclear layer, INL – inner nuclear layer, IPL – inner plexiform layer, GCL – ganglion cell layer.

Since components of the NF-κB pathway are expressed by both Müller glia and microglia (Fig. 1), pharmacological manipulations of the pathway are expected to directly influence both cell types. Thus, we investigated how NF-κB-signaling influences Müller glia reprogramming in the absence of microglia. In the absence of microglia, inhibition of NF-κB-signaling with sulfasalazine or PGJ2 following NMDA-treatment had no significant effect upon numbers of proliferating MGPCs (Figs. 7d,e,g), or on total numbers of TUNEL^+^ dying cells (Figs. 7f,h). Treatment with SC757 produced the same outcomes as sulfasalazine and PGJ2 treatment (not shown). Taken together, these findings suggest that in the absence of microglia, NF-κB-inhibitors have no effect because there was little or no signaling to inhibit. It is possible that activated microglia provide the signals required to activate NF-κB in Müller glia after damage.

We next investigated whether activation of NF-κB in the absence of microglia influenced the formation of MGPCs. In retinas treated with clodronate liposomes, a high dose (1 µmol) NMDA was applied in combination with prostratin, TNFSF15, or vehicle controls. Interestingly, treatment with prostratin or TNFSF15 caused significant increases in numbers Sox2^+^/EdU^+^ cells in microglia-depleted, NMDA-damaged retinas (Figs. 7i-k). These data suggest that in the absence of microglia, activation of NF-κB promotes MGPC-proliferation, which is opposite to the effects of NF-κB activation on MGPC-proliferation in damaged retinas with reactive microglia (Fig. 4). Additionally, in the absence of microglia, treatment of damaged retinas with TNFSF15 significantly increased numbers of TUNEL^+^ cells (Figs. 7m,n), whereas prostratin had no effect (Fig. 7l). Collectively, these findings suggest that reactive microglia are required to induce NF-κB activation in Müller glia and initiate a gliotic response which progresses into reprogramming, but sustained activation of NF-κB suppresses proliferation of MGPCs.

### Expression patterns of NF-κB-related genes in retinal cells after treatment with insulin and FGF2

In the chick retina, MGPCs are known to form in the absence of retinal damage in response to 3 consecutive daily injections of insulin and FGF2 (Fischer et al., 2002). Accordingly, we establish scRNA-seq libraries for retinas treated with insulin and FGF2 (Fig. 8a). Müller glia were identified based on combined expression of *Vim, Sox2* and *Sox9*, and MGPCs were identified based on down-regulation of *Glul* and *Rlbp1*, and up-regulation of *Pcna, Cdk1* and *Nestin* (Fig. 8b). We analyzed expression of NF-kB components in Müller glia and MGPCs, since treatment with insulin and FGF2 was not expected to have significant effects upon retinal neurons. *Nfkbia* had scattered expression in Müller glia and MGPCs, and appeared to be expressed in reduced numbers of non-proliferative Müller glia treated with 3 doses of insulin and FGF2 (Fig. 8c). By comparison, *Nfkbib* was expressed by relatively high numbers of Müller glia in control and by MGPCs, but in diminished numbers of non-proliferative Müller glia treated with insulin and FGF2 (Fig. 8c). *Nfkbiz* was expressed by Müller glia in control retinas, and appeared to be down-regulated in Müller glia and MGPCs treated with insulin and FGF2 (Fig. 8c). *Tnfsf15* was detected only in microglia (Fig. 8c). By comparison, *Tnfsf10* was detected in relatively few Müller glia treated with saline or 2 doses insulin and FGF2, but was expressed in many Müller glia and MGPCs treated with 3 doses of insulin and FGF2 (Fig. 8c). *Tnfrsf1a* was detected at relatively high levels in scattered Müller glia in control and growth factor-treated retinas (Fig. 8c). By comparison, *Tnfrsf21* was widespread in control Müller glia and was down-regulated in Müller glia from growth factor-treated retinas, and modestly decreased in MGPCs compared to resting Müller glia (Fig. 8c).

**Figure 8:**
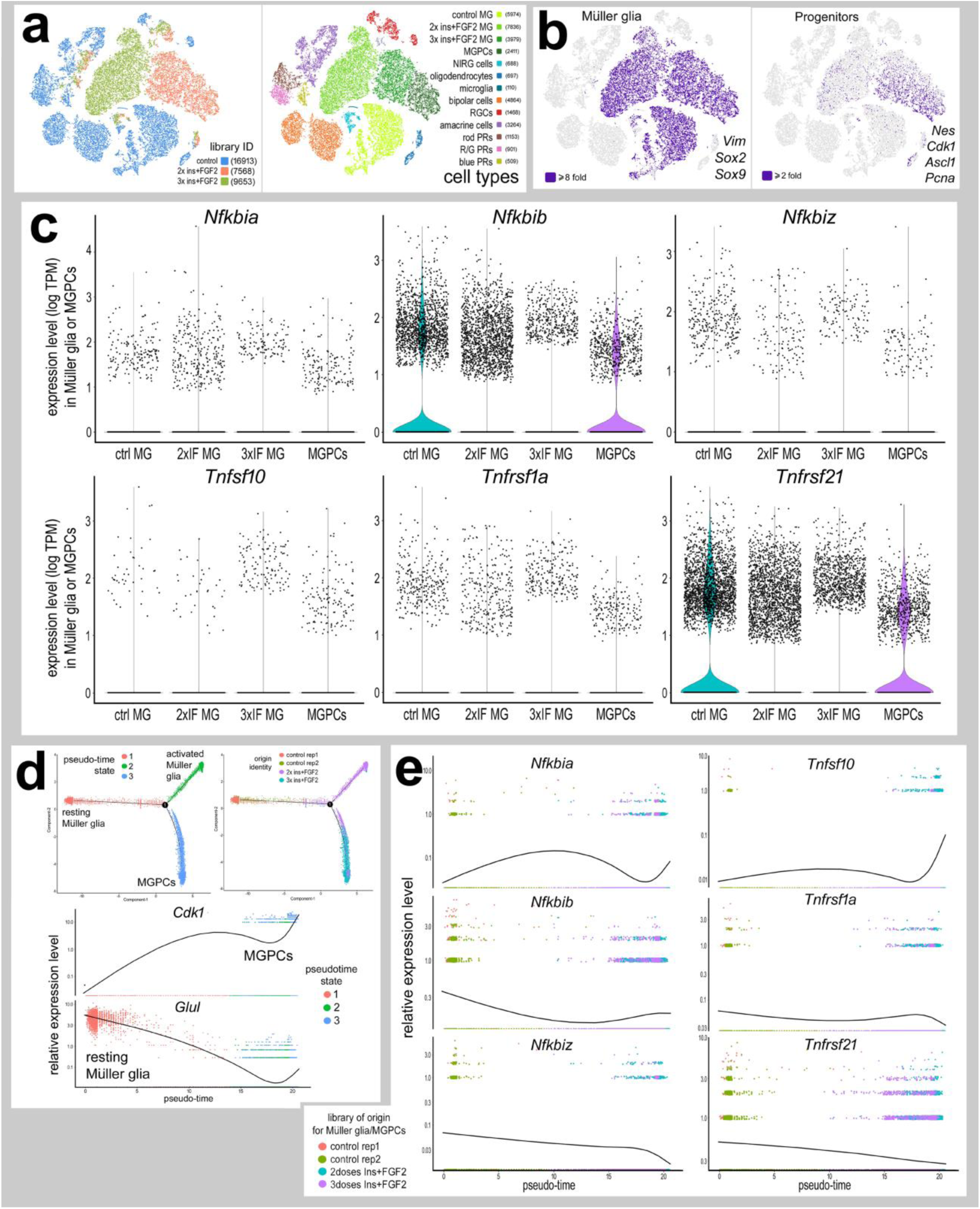
Expression patterns of NF-κB-related genes in retinal cells after treatment with insulin and FGF2. Libraries for scRNA-seq were established for cells from control retinas and from retinas at treated with 2 or 3 consecutive daily doses of insulin and FGF2. Each dot represents one cell. Cells were sampled from control retinas (16913 cells), and from retinas treated with 2x doses (7568 cells) or 3x doses (9653 cells) (**a**). tSNE plots show distinct clustering of different retinal cell populations and numbers of cells surveyed within each cluster (in parentheses) (**a**). Each individual point represents one cell. Müller glia were identified based on expression of *Sox2, Sox9*, and *Vimentin* (**b**). Progenitors were identified based on expression of *Cdk1, Ascl1, PCNA*, and *Nestin* (**b**). Violin/scatter plots for expression levels of *Nfkbia, Nfkbib, Nfkbiz, Tnfsf10, Tnfrsf1a* and *Tnfrsf21* in Müller glia in control retinas, 2x insulin and FGF2, 3x insulin and FGF2 and MGPCs (**c**). Seurat objects identified as Müller glia and MGPCs were re-embedded for pseudotime analysis. Monocle was used to generate pseudotime trajectories in an unbiased manner guided by the differentially expressed genes. Pseudotime states included: (1; peach) resting Müller glia, (2; olive) transitional MG, (3; green), and (3; blue) MGPCs (**d**). Reduction of pseudotime to the X-axis placed the resting Müller glia to the far left and the MGPCs to the far right (**d**). Relative levels of expression among Müller glia and MGPCs across pseudotime was assessed for components of the NFkB-pathway; *Nfkbia, Nfkbib* and *Nfkbiz*, and TNF-related factors and receptors; *Tnfsf10, Tnfrsf1a* and *Tnfrsf21* (**e**).

Pseudotime analysis, using an unbiased comparison of highly variable genes, revealed 3 distinct states and a trajectory that includes distinct branches for resting Müller glia, 2 doses insulin+FGF2 (IF) and 3 doses IF (Fig. 8d). Resting Müller glia, from control retinas were located to the left of the pseudotime axis, and expressed high levels of mature glial markers, such as *Glul* (Fig. 8d). MGPCs were located to the far right of the pseudotime axis, and expressed high levels markers for proliferation and progenitors, such as *Cdk1* (Fig. 8e). *Nfkbia* and *Tnfsf10* were low in resting Müller glia, increased in transitional glia, decreased in activated glia, but increased in MGPCs (Fig. 8e). Similar to trends seen in damaged retinas, levels of *Nfkbib* and *Nfkbiz* were relatively high in resting MG, decreased in transitional glia, and *Nfkbib* increased while *Nfkbiz* decreased toward (Fig. 8e). Relative expression of *Tnfrsf1a* and *Tnfrsf21* was high in resting Müller glia, decreased in transitional, and further decreased in MGPCs (Fig. 8e).

### NF-κB signaling influences MGPC formation in the absence of retinal damage

We next investigated whether NF-κB-signaling influences the formation of MGPCs in the absence of retinal damage. We found that 4 consecutive daily injections of sulfasalazine, prostratin, or TNFSF15 alone failed to induce proliferation of MGPCs and did not induce cell death (data not shown). We next tested whether the combination of Fibroblast growth factor (FGF2) and sulfasalazine, prostratin, or TNFSF15 influenced the formation of proliferating MGPCs. Four doses of FGF2 alone is able to activate a network of signaling pathways that promotes the formation of proliferating MGPCs in the absence of damage in the chick retina (Fischer et al., 2009a; Gallina et al., 2015; Todd et al., 2016; Zelinka et al., 2016). Four consecutive daily treatments with NF-κB inhibitors (sulfasalazine, PGJ2, or SC757) in combination with FGF2 significantly increased the numbers of EdU^+^/Sox9^+^ proliferating MGPCs compared to treatment with FGF2 alone (Fig. 9a-c). Additionally, application of sulfasalazine in FGF2-treated retinas resulted in decreased microglia reactivity, indicated by decreased levels CD45-immunofluorescence (Fig. 9d,e). Application of NF-κB activator, prostratin, in FGF2-treated retinas resulted in a significant decrease in the number of EdU^+^/Sox9^+^ proliferating MGPCs (Figs. 9f,g), but did not influence the proliferation of microglia (Fig. 9h). Similarly, treatment with TNFSF15 and FGF2 resulted in a significant decrease in MGPC proliferation compared to FGF2 alone, while there was no change in microglial proliferation (Figs. 9g,h).

**Figure 9:**
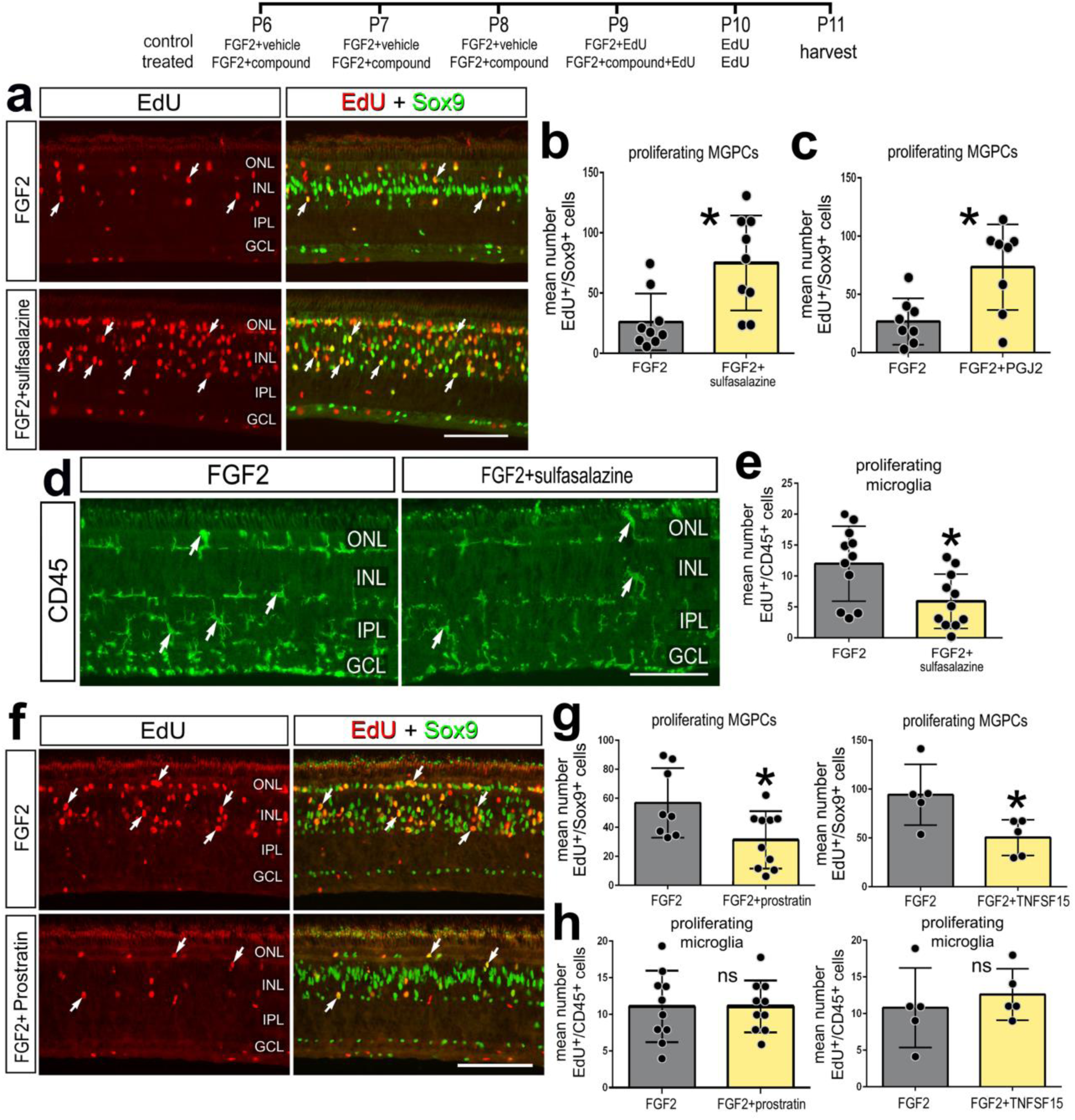
Inhibition of NF-κB promotes MGPC proliferation in the absence of damage. Retinas were treated with FGF2 plus NF-κB inhibitors (sulfasalazine or PGJ2) or NF-κB activators (prostratin or TNFSF15) or vehicle controls at P6-P9, EdU was added to injections at P8-P10, and retinas harvested at P11. Retinal sections were labeled for EdU (red; **a,f**), or with antibodies to Sox9 (green; **a,f**), or CD45 (green; **d**). Histograms in **b**, **c** and **g** represent mean number (± SD and individual data points) of proliferating MGPCs. Histograms in **e** and **h** represents mean number (± SD and individual data points) of proliferating microglia. Arrows represent proliferating MGPCs (**a,f**) or microglia (**d**). Calibration bars in **d, i,** and **n** represent 50 µm. Abbreviations: ONL – outer nuclear layer, INL – inner nuclear layer, IPL – inner plexiform layer, GCL – ganglion cell layer.

## Discussion

Our findings indicate that components of the NF-κB pathway are dynamically expressed in Müller glia during the process of reprogramming into MGPCs. Further, our findings indicate that active NF-κB-signaling in damaged retinas acts to suppress the formation of proliferating MGPCs and may contribute to detrimental inflammation that exacerbates cell death. Interestingly, the effects of activation of NF-κB on the formation of MGPCs in damaged retinas are reversed when the microglia are absent. We propose that pro-inflammatory signals from microglia are required to induce NF-κB activation and initiate a gliotic response which progresses into reprogramming, but this reprogramming is suppressed by sustained activation of NF-κB. Finally, we find that NF-κB is part of the network of pathways activated by FGF2-treatment, and in the absence of neuronal damage NF-κB activation suppresses the formation of MGPCs. These findings are summarized in Figure 10.

**Figure 10:**
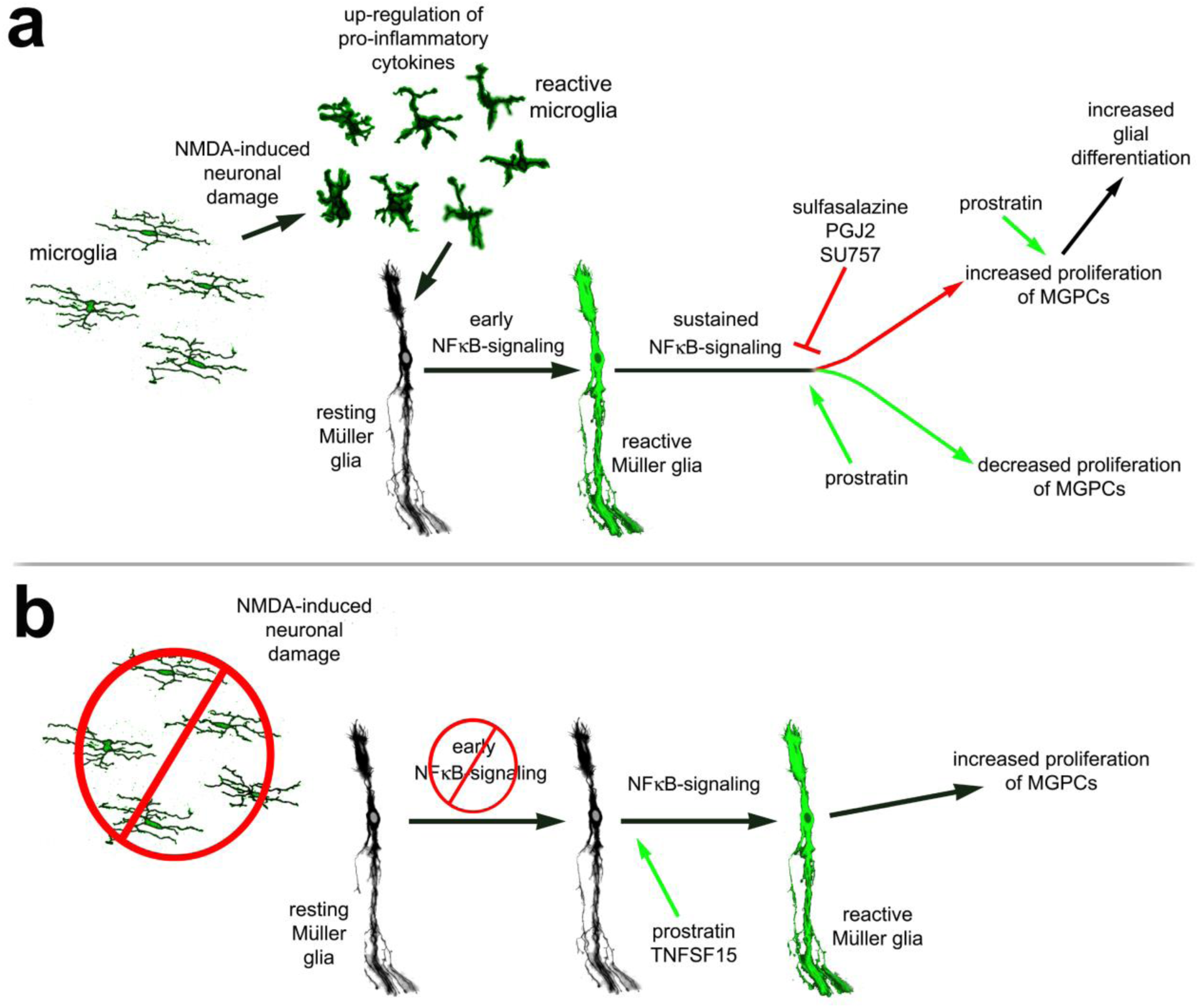
Schematic summary of findings. **(a)** After NMDA, microglia are required for activation of NF-κB signaling. Inhibition of NF-κB promotes Müller glia reprogramming into proliferating MGPCs, while sustained activation suppresses the formation of MGPCs. NF-κB activation in the progeny of MGPCs promotes glial fate. **(b)** In the absence of microglia, NF-kB signaling is diminished in Müller glia, and MGPC formation is not initiated. Stimulation of NF-κB signaling after damage promotes Müller glia reactivity to initiate reprogramming and promote the formation of proliferating MGPCs.

Our findings are consistent with the notion that pro-inflammatory signals from reactive microglia activate NF-κB in Müller glia in damaged retinas. Microglia become highly reactive in NMDA-damaged retinas (Fischer et al., 1998; Fischer et al., 2014; Todd et al., 2019; Wada et al., 2013), and participate in bi-directional communication with Müller glia (Wang et al., 2011; Wang et al., 2014). Ablation of microglia prior to retinal damage suppresses the formation of MGPCs in the chick retina (Fischer et al., 2014). In NMDA-damaged retinas, microglia rapidly and transiently up-regulate IL-1α, IL-1β and TNFα (Todd et al., 2019), and these cytokines are known to activate NF-κB in different cells and contexts (Hayden and Ghosh, 2004; Osborn et al., 1989). In most instances the formation of MGPCs requires neuronal damage, and levels of neuronal damaged are positively correlated to numbers of proliferating MGPCs (Gallina et al., 2014b; Todd et al., 2017). However, there are examples where proliferating MGPCs form in the absence neuronal death in chick and fish model systems (Fischer et al., 2002; Fischer et al., 2009a, 20014; Wan et al., 2012; Wan et al., 2014). Despite decreased cell death in NMDA-damaged retinas treated with NF-κB-inhibitors, we find increased MGPC-proliferation with reactive microglia present, whereas there was no change in MGPC-proliferation in damaged retinas treated with NF-κB inhibitors when microglia were absent. Our findings suggest that the relationship between levels of damage and formation of MGPCs can be uncoupled by the inhibition of NF-κB. Similarly, the application of FGF2 prior to NMDA-induced damage is potently neuroprotective and facilitates the formation of MGPCs (Fischer et al., 2009a). Even in the absence of neuronal damage reactive microglia are required for the formation of proliferating MGPCs (Fischer et al., 2014). Collectively, these findings suggest that pro-inflammatory signals provided by microglia are required to initiate Müller glial reactivity as a first step toward reprogramming into MGPCs.

NF-κB-signaling is coordinated with a network of cell-signaling pathways that regulate the reprogramming of Müller glia into MGPCs. Many different cell-signaling pathways are known to participate in the formation of MGPCs in the chick retina, including MAPK (Fischer et al., 2009a; Fischer et al., 2009b), Jak/Stat (Todd et al., 2016; Zhao et al., 2014), Notch (Ghai et al., 2010; Hayes et al., 2007), Hedgehog (Todd and Fischer, 2015; Wan et al., 2007), glucocorticoid (Gallina et al., 2014b), retinoic acid (Todd et al., 2018), BMP/TGFβ (Todd et al., 2017), and Wnt/β-catenin(Gallina et al., 2015). There is evidence for cross-talk between many of these pathways with NF-ĸB signaling in different cellular contexts (Ahmad et al. 2015; Ma and Hottiger 2016; Mohan et al. 1998; Nelson et al. 2013). For example, there is evidence of bidirectional communication between Wnt and NF-ĸB signaling, and we have previously shown that Wnt-signaling promotes formation of proliferating MGPCs in the chick retina (Gallina et al., 2015), which is consistent with findings in mouse and fish retinas (Meyers et al., 2012; Meyers et al., 2012; Osakada et al., 2007; Yao et al., 2016; Yao et al., 2018). It has been shown that over-expression of β-catenin, a transcriptional effector of Wnt, suppresses NF-ĸB activity and expression of NF-ĸB target genes (reviewed by (Ma and Hottiger, 2016). Conversely, NF-ĸB activity can suppress Wnt-signaling by directly inhibiting β-catenin or indirectly via activating GSK3β (Ma and Hottiger, 2016). These data are consistent with our results showing that inhibition of NF-κB results in increased proliferation of MGPCs and previous work showing activation of Wnt-signaling promotes the proliferation of MGPCs (Gallina et al., 2015). Additional studies are required to further understand the relationship between these two signaling cascades during Müller glia reprogramming.

It is possible that some of the reprogramming-suppressing effects of NF-κB-signaling are mediated by Smad2/3. *In vitro*, NF-ĸB can induce cell cycle arrest and terminal differentiation via IKKα-dependent regulation of Smad2/3 target genes (Descargues et al., 2008). We have reported that TGFβ/Smad2/3-signaling inhibits, whereas BMP4/Smad1/5/8-signaling promotes the formation of proliferating MGPCs in the chick retina (Todd et al., 2017). By contrast, the influence of NF-κB on the proliferation of MGPCs may not depend on Notch. In the chick retina, inhibition of Notch-signaling with gamma-secretase inhibitor, DAPT, suppresses the proliferation of MGPCs (Ghai et al., 2010; Hayes et al., 2007). We found that DAPT failed to suppress MGPC proliferation in damaged retinas treated with NF-κB inhibitor, suggesting that NF-κB takes precedence over or acts downstream and overrides of influence of Notch on the proliferation of MGPCs. Further studies are required to better establish where NF-κB fits into the hierarchy of cell signaling pathways that regulate the formation of MGPCs.

Differences in activation of NF-κB may underlie differences in the reprogramming potential of MGPCs in different vertebrates. In the mouse retina, it has been shown that NMDA-induced excitotoxic damage activates NF-κB-signaling in Müller glia (Lebrun-Julien et al., 2009). In zebrafish retina, TNFα is required for MGPC proliferation via activation of Stat3 (Nelson et al., 2013). A recent comparative transcriptomic and epigenomic study has indicated that components of the NF-κB pathway are prominently expressed and regulatory elements accessible for transcription in mouse Müller glia rapidly following NMDA-induced damage (Hoang et al., 2019 preprint). By comparison, Müller glia in the chick retina modestly up-regulate relatively few components of the NF-κB-pathway, and Müller glia in the zebrafish do not express nor dynamically regulate components of the NF-κB-pathway (Hoang et al., 2019 preprint). Collectively, the study by Hoang and colleagues (2019) suggest that the reprogramming of Müller glia begins with acquisition of a reactive, activated phenotype then progression through reactivity, down-regulation of glial genes and acquisition of progenitor phenotype and proliferation; this occurs for the Müller glia in fish and chick retinas, but not for the Müller glia in mouse retinas. Rodent Müller glia rapidly transition into an activated state and then revert back to a resting state (Hoang et al., 2019 preprint). Consistent with this notion, Thomas and colleagues (2016) provide data to suggest that Müller glia in light-damaged zebrafish retina acquire a reactive, neuroprotective phenotype prior to transitioning through to reprogramming into a proliferating, progenitor phenotype (Thomas et al., 2016). Collectively, these data suggest the following model for the role of NF-κB in reprogramming of Müller glia into MGPCs (Fig. 10); (1) neuronal damage rapidly induces microglial reactivity, (2) increased production of pro-inflammatory cytokines from microglia, (3) a rapid, initial activation of NF-κB in Müller glia that initiates gliosis, (4) transition to reprogramming (down-regulation of glial genes and up-regulation of progenitor genes), but (5) sustained NF-κB maintains reactive phenotype and pushes glia back to a resting phenotype (mouse) or suppresses reprogramming of Müller glia into proliferating MGPCs (chick). In the absence of reactive microglia, our data suggest that the process of reprogramming requires initiation of glial reactivity and de-differentiation, which can be provided by exogenous TNFSF15 or NF-κB-agonist.

We found that activation of NF-κB promoted the differentiation of Müller glia from the progeny of proliferating MGPCs. Our findings are consistent with reports that pro-inflammatory cytokines and NF-κB promote gliogenesis during neural development (Bonni et al., 1997; Deverman and Patterson, 2009; Mondal et al., 2004). For example, IL-1 promotes acquisition of astrocyte cell fate, with peak expression concurrent with the generation of astrocytes in development (Giulian et al., 1988). Further, IL-1 is known to activate NF-κB-signaling and there is evidence that NF-ĸB promotes specification of astrocytes (Mondal et al., 2004). Additionally, TNFα stimulates hippocampal neural precursors to promote astrocyte formation, and this occurs via up-regulation of the pro-glial/progenitor gene bHLH transcription factor Hes1 (Keohane et al., 2010). Collectively, these findings suggest that NF-κB acts to promote glial differentiation from MGPC progeny.

NF-κB influences cell death in a context-specific manner. NF-κB has been shown to promote neuron death (Schneider et al., 1999), but has also been shown to support neuronal survival (Bhakar et al., 2002). NF-κB signaling is activated by a variety of factors including TNFα, reactive oxygen species, lipopolysaccharide, and various growth factors (Schreck et al., 1991). In the mouse retina, NF-κB signaling has been shown to be involved in NMDA-induced retinal neuron death, and inhibition of NF-κB signaling prevents retinal neuron death (Lebrun-Julien et al., 2009). Consistent with these findings, we found that inhibition of NF-κB reduced numbers of dying amacrine and bipolar neurons in NMDA-damaged retinas and promoted the survival of ganglion cells in colchicine-damaged retinas. This may have resulted from suppressed production of TNF-ligands from reactive microglia.

## Conclusions

Our findings suggest that NF-κB-signaling plays significant roles in regulating glial reactivity, neuronal survival, and the reprogramming of Müller glia into proliferating MGPCs. NF-κB-signaling activity suppresses the formation of proliferating MGPCs, and the influence of NF-κB depends on reactive microglia. Despite reducing levels of cell death, NF-κB-inhibitors also stimulated the formation of MGPCs. In the absence of damage, in retinas treated with FGF2, NF-κB-signaling is recruited into a network of cell-signaling pathways that regulate the reprogramming of Müller glia into proliferating MGPCs. We propose that reactive microglia may provide signals to activate Müller glia via NF-κB to initiate the process of reprogramming.

## Acknowledgements

This work was supported by grants from NIH (NEI RO1 EY022030-06 to AJF; UO1 EY027267-03 to AJF and SB). Thanks to Alex Campbell and Levi Todd for comments and discussions that shaped the final form of the paper.

## Author Contributions

I.P. designed and executed experiments, gathered data, constructed figures and contributed to writing the manuscript; K.D. executed experiments, gathered data, and constructed figures; S.B. and T.H. contributed to creation of sc-RNA seq libraries; A.J.F. designed experiments, constructed figures and contributed to writing the manuscript. All authors read and approved final manuscript.

## Supplemental Data

**Supplemental Figure 1:**
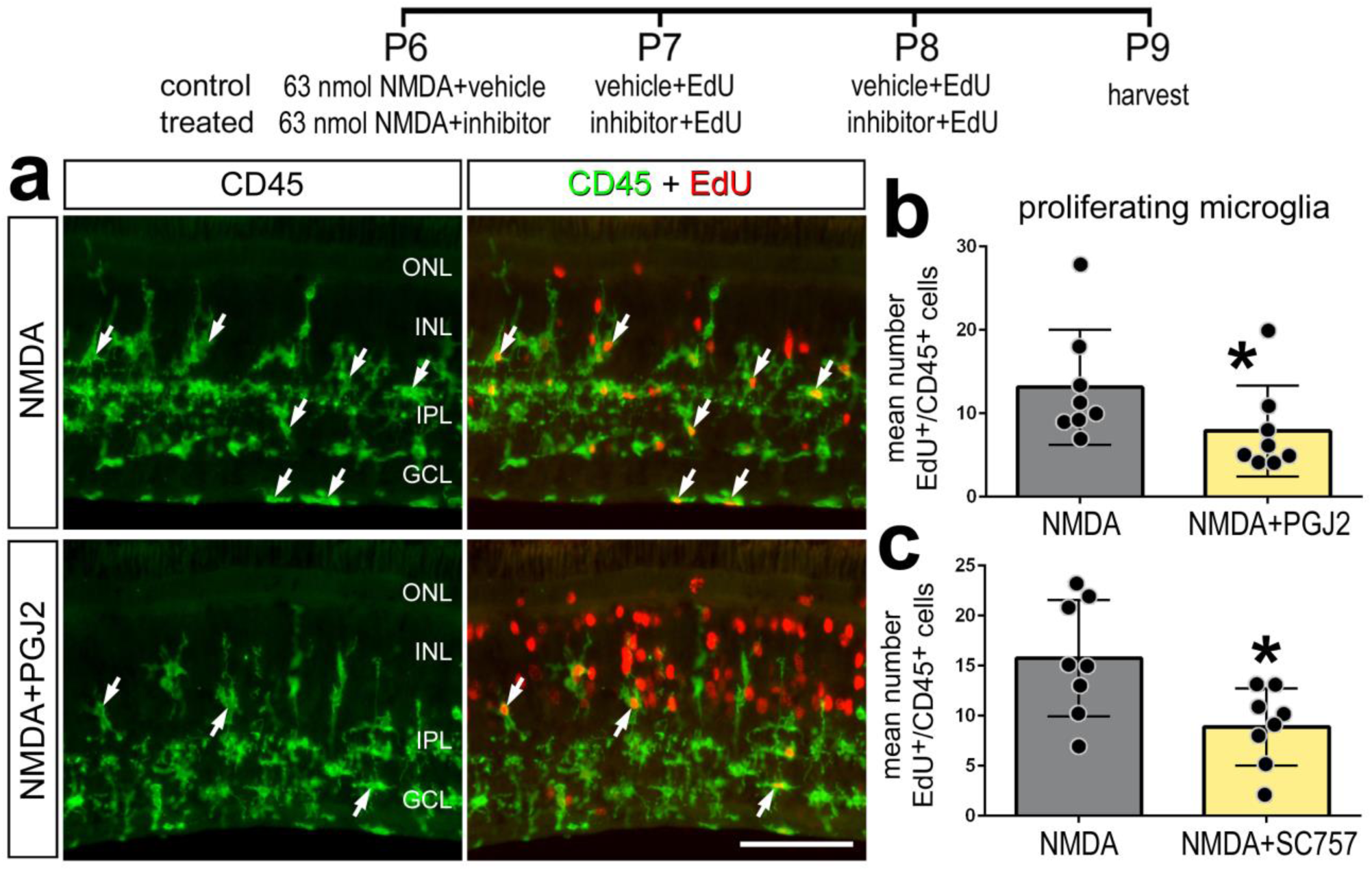
Inhibition of NF-κB after damage results in decreased microglia proliferation. Retinas were treated with NMDA plus vehicle or NMDA plus NF-κB inhibitors (PGJ2 or SC757) at P6-P8. Edu was added to injections at P7-P8, and retinas were harvested at P9. Retinal sections were labeled for CD45 (green; **a**) and Edu (red; **a**). Histograms in **b** and **c** represent mean number (± SD and individual data points) of proliferating microglia. Arrows represent proliferating microglia in **a**. Calibration bars in **a** represent 50 µm. Abbreviations: ONL – outer nuclear layer, INL – inner nuclear layer, IPL – inner plexiform layer, GCL – ganglion cell layer.

**Supplemental Figure 2:**
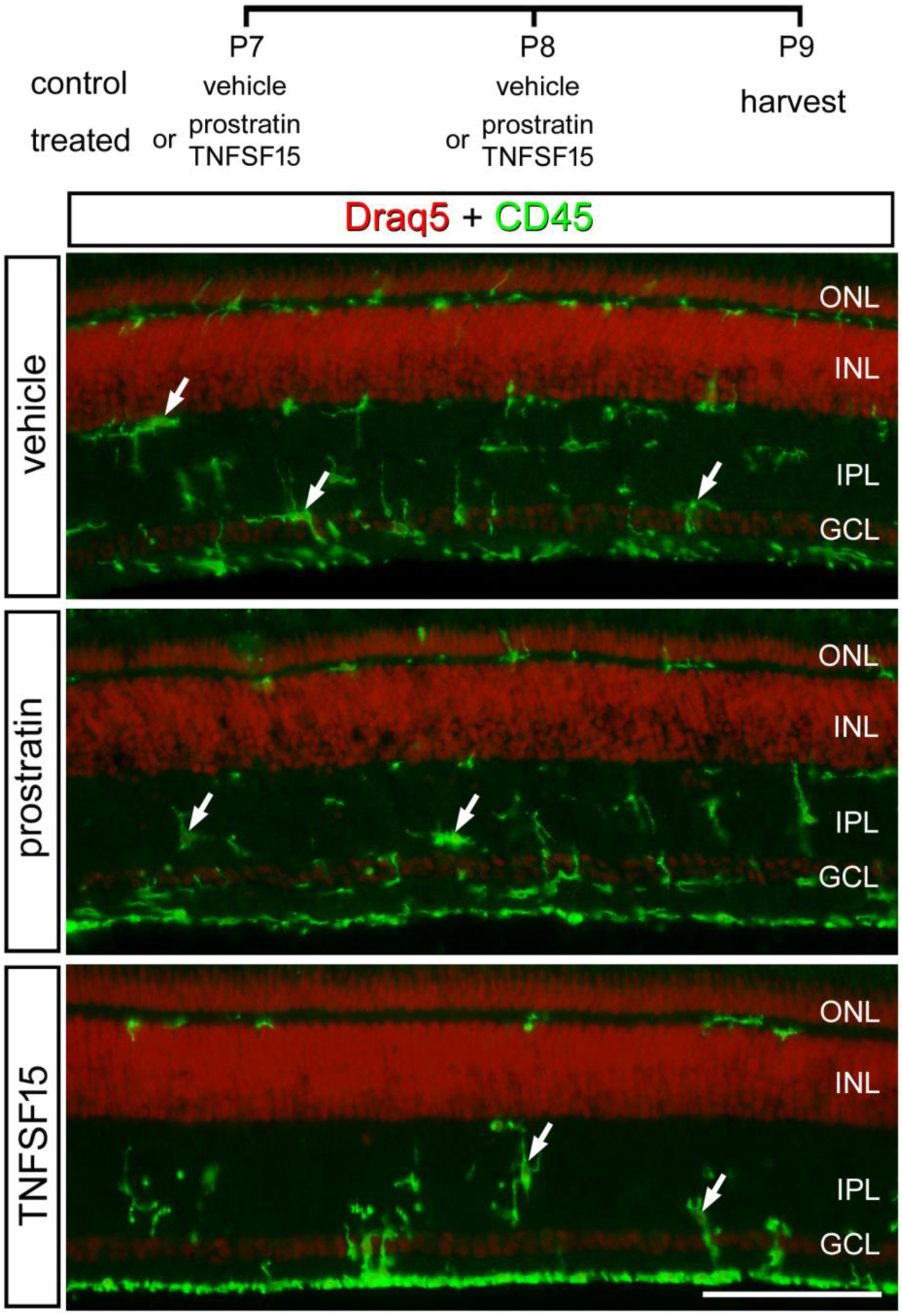
Microglia in retinas treated with prostratin or TNFSF15. Eyes were treated with vehicle, prostratin or TNFSF15 at P7 and P8, and retinas harvested at P9. Sections of the retina were labeled with DRAQ5 (red nuclei) and antibodies to CD45 (green). Arrows indicate microglia. The calibration bar represents 50 µm. Abbreviations: ONL – outer nuclear layer, INL – inner nuclear layer, IPL – inner plexiform layer, GCL – ganglion cell layer.

